# HDMAX2-surv: high-dimensional mediation analysis of survival data with application to pancreatic cancer

**DOI:** 10.1101/2025.09.09.675033

**Authors:** Florence Pittion, Elise Amblard, Emilie Devijver, Adeline Samson, Nelle Varoquaux, Magali Richard

## Abstract

**Motivation:** High-dimensional mediation analysis is critical for dissecting causal pathways in complex diseases. However, existing methods struggle to handle censored survival outcomes and unobserved confounders in epigenetic data. There is a pressing need for scalable, statistically robust frameworks to identify epigenetic mediators of exposure–outcome relationships, especially in cancer research where survival data are central and molecular pathways are highly complex and interconnected.

**Results:** We introduce HDMAX2-surv, a two-step framework extending HDMAX2 to survival analysis. HDMAX2-surv integrates (i) latent factor modeling to adjust for unobserved confounders, (ii) flexible survival models (Aalen additive hazards and accelerated failure-time models). Simulations demonstrated superior performance over state-of-the-art methods (e.g., HIMA) in mediator selection and effect estimation. We integrated this approach with causal discovery frameworks and immune deconvolution algorithms to dissect multi-pathway mediation mechanisms. Applied to TCGA pancreatic adenocarcinoma data (n=112), HDMAX2-surv identified 36 aggregated methylated regions (AMRs) mediating tobacco exposure effects on survival, including immune-mediated pathways undetectable via gene expression alone.

**Availability and implementation:** HDMAX2-surv is implemented in R and available on GitHub (https://github.com/bcm-uga/tims-pdac), with documentation and example scripts for reproducibility.

## Introduction

Understanding the causal mechanisms linking environmental exposures to clinical outcomes remains a central challenge in biomedical research. Mediation analysis offers a comprehensive framework to identify molecular mediators, such as epigenetic modifications, that underlie exposure–outcome pathways (1; 2). However, extending mediation analysis to high-dimensional omics data in the presence of censored survival outcomes introduces substantial statistical and computational challenges. These include: (i) scalable mediator selection strategies (e.g., Sure Independence Screening (SIS), False Discovery Rate (FDR) control, or regularization approaches (3)); (ii) appropriate specification of the survival data, whether non-parametric (Kaplan-Meier or Nelson-Aalen estimators), parametric (via Weibull, log-normal, exponential distributions), or semi-parametric (Cox proportional hazards models, 4; 5); and (iii) the integration of these components into a coherent mediation framework (e.g., Sobel test (6), Baron and Kenny method (7), bootstrapping, sensitivity analysis (1; 2)).

Several statistical methods have been proposed to address these challenges. HIMA combines sure independence screening (SIS) for the exposure–mediator model with minimax-concave-penalty (MCP) regularisation in the Cox outcome model to construct a sparse mediation set (8; 9). MASH couples SIS and MCP variable selection and utilizes *R*^2^-based measures to estimate mediation effects within the Cox regression model (10). IUSMMT proposes a Cox model with random effects to identify mediators in high-dimensional settings (11). CoxMKF uses aggregation of multiple knockoffs with Cox proportional hazards models to control the FDR (12). mediateR estimates high-dimensional mediation via dual ridge penalties and joint significance testing, and handles continuous, binary, and survival outcomes (13). Finally, SMAHP employs a multi-omic methodology capable of handling both high-dimensional mediators and exposures with FDR control (14). Despite these advances, no consensus exists on the optimal methodological approach for high-dimensional mediation analysis in survival studies.

To overcome these limitations, we introduce HDMAX2-surv, a two-step framework extending HDMAX2 (15) to censored survival outcomes. HDMAX2-surv integrates: (i) a latent factor modeling to adjust for unobserved confounders in high-dimensional molecular data; (ii) a high-dimensional mediation significance test with controlled FDR; and (iii) a flexible survival modeling using both Aalen additive hazards and accelerated failure-time models, thereby relaxing proportional hazards assumptions. In contrast to the original HDMAX2 approach (16), which was not designed for time-to-event outcomes, HDMAX2-surv explicitly models censored survival data.

We validate HDMAX2-surv thoroughly using simulated data and demonstrate its utility in pancreatic ductal adenocarcinoma (PDAC), a highly lethal malignancy accounting for approximately 90% of pancreatic cancers and ranking among the leading causes of cancer-related mortality in Europe (17). Tobacco exposure is a well-established risk factor for both PDAC incidence and prognosis (18; 19; 20), with meta-analyses reporting a 75% increased in incidence risk among smokers (19). Despite this strong epidemiological evidence, the molecular mechanisms linking tobacco exposure to PDAC survival, particularly through epigenetic regulation and immune-related processes, remain insufficiently characterized (21). Applying HDMAX2-surv to PDAC samples from The Cancer Genome Atlas (TCGA; *n* = 112), we identified 36 aggregated methylated regions (AMRs) mediating the effect of tobacco exposure on overall survival. Integrating causal discovery and serial mediation analysis further revealed multi-step pathways involving immune-related mechanisms that were not detectable using gene expression data alone.

Together, these results establish HDMAX2-surv as a scalable and statistically robust framework for high-dimensional mediation analysis with survival outcomes, applicable to complex diseases where omics mediators and time-to-event endpoints intersect.

## Method

HDMAX2-surv is a high-dimensional mediation analysis framework specifically designed for censored survival outcomes. The method follows the directed acyclic graph illustrated in Figure 1A, where we evaluate the causal effect of exposure (*X*) on survival outcome (*S*), and quantify mediation (*M*) in a high-dimensional setting. Both observed (*O*) and unobserved (*U*) confounding factors are accounted for through a latent factor modeling strategy.

**Figure 1.**
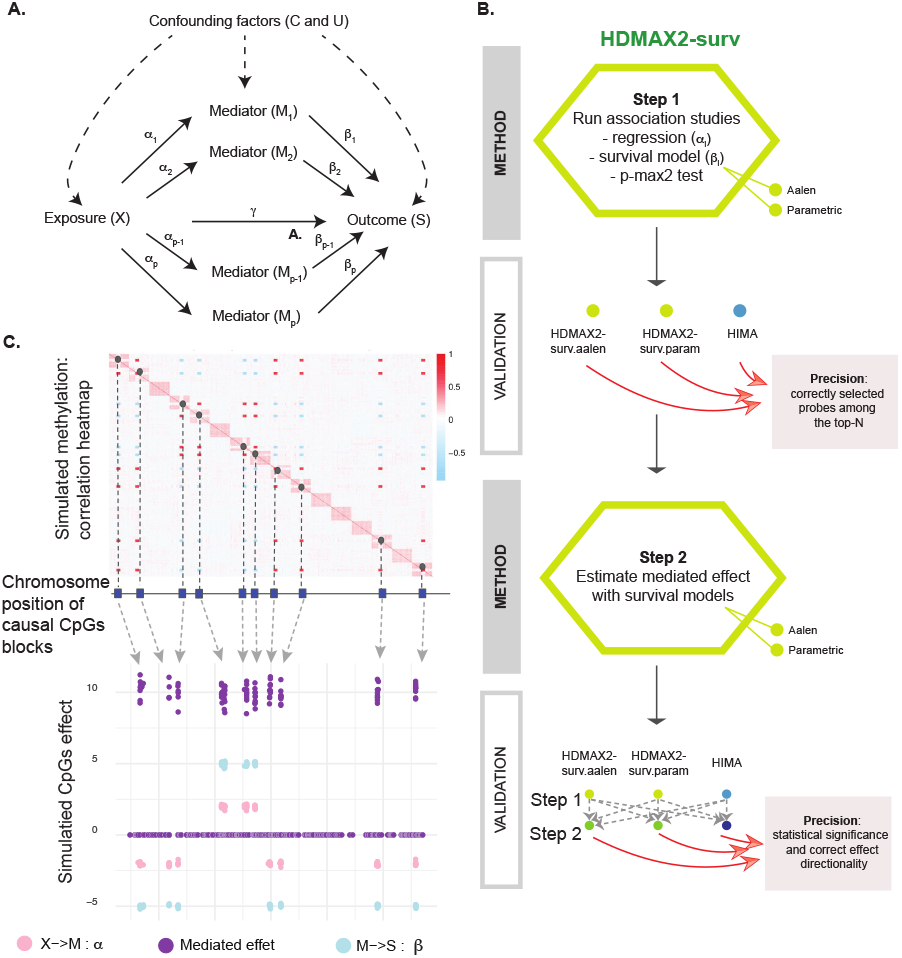
HDMAX2-surv: theoretical model, algorithm, and validation design. **(A)** Directed acyclic graph describing the high-dimensional mediation framework with an exposure *X, p* potential mediators *M*, a survival outcome *S*, and confounding factors (observed *C* and unobserved *U*). **(B)** Schematic representation of the HDMAX2-surv method (Step 1: selection of candidate mediators; Step 2: mediation effect estimation), with a two-step evaluation workflow based on estimation precision using simulated data with known ground truth. `HDMAX2-surv.aalen` and `HDMAX2-surv.param` were compared to `HIMA`, a method designed for high-dimensional survival mediation analysis. **(C)** Schematic representation of the simulation framework. Heatmap shows pairwise correlations among simulated DNAm probes (CpGs), illustrating the dependence structure imposed in the simulated data (blocks of correlated CpGs). We simulated genomic positions on a virtual chromosome. For selected blocks of causal CpGs (dark blue squares), corresponding to true mediators in our simulations, we assigned positive mediation effects with identical effect values for all CpGs within each block (purple). As we used Cox models to simulate survival outcomes, positive mediated effects are interpreted as deleterious for survival. This mediated effect represents the product of simulated *α* effects (*X* → *M*, pink) and *β* effects (*M* → *S*, blue).

HDMAX2-surv extends the original HDMAX2 method (16; 15) to accommodate time-to-event outcomes. The original HDMAX2 framework relies on a latent factor mixed regression model (LFMM; Equations 1 and 2) and performs high-dimensional association testing in which statistical significance of candidate mediators is assessed using a max-squared test (Equation 3). Mediation effect is then estimated by causal mediation analysis, within the counterfactual framework, using the `mediation` R package (2) which estimates the Average Causal Mediation Effect (ACME) from mediator model *M* (Equation 1) and continuous or binary outcome model *Y* (Equation 2).

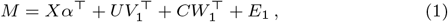

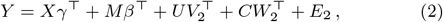

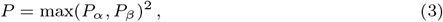

with *M*: the mediator matrix, *X*: the exposition matrix, *Y*: the outcome (continuous or binary), *C*: the observed confouding factors, *U*: the estimated score matrix for the unobserved latent factors, *E*: the residual noise. A significance value *P*_*α,j*_ is computed by the HDMAX2 method for the test of a null effect size for exposure variable *X* on intermediary variable *M*_*j*_, for each *j*, as well as a significance value *P*_*β,j*_ corresponding to the association of each intermediary variable *M*_*j*_ with the outcome variable *Y*.

A central methodological challenge addressed in HDMAX2-surv is the integration of censored survival outcomes within a high-dimensional mediation framework. Cox proportional hazards models are widely used in survival analysis because they do not require specification of the baseline hazard function. However, these models rely on the proportional hazards assumption and yield non-collapsible hazard ratios, meaning that inclusion of a mediator may alter the estimated hazard ratio even in the absence of true mediation. Consequently, coefficients from models with and without the mediator are not directly comparable, complicating indirect effect estimation. In contrast, parametric survival models and additive hazard models do not exhibit this limitation (4). Following the recommendations of Lapointe-Shaw *et al*. (4), we therefore implemented two alternative survival modeling strategies (Equations 4 and 5) within the HDMAX2-surv framework (Figure 1B), replacing the original regression-based outcome model for *Y* used in HDMAX2 (Equation 2).

The first approach relies on a parametric accelerated failure time (AFT) model:

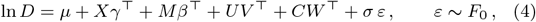

where *D* denotes the death time with a survival function *S*(*t*), *µ* is the intercept, and *σ* is a scale parameter. The error distribution *F*_0_ determines the parametric form of the survival model: (i) *F*_0_ = EV1, the extreme value type 1 distribution, yields the Weibull model, (ii) *F*_0_ = Logistic yields the log-logistic model, and (iii) *F*_0_ = *N* (0, 1) yields the log-normal model. The best-fitting distribution for the censored data was selected using the Akaike Information Criterion.

The second approach is based on the Aalen additive hazards model:

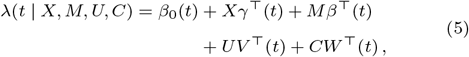

where *λ*(*t* | *X, M, U, C*) denotes the hazard rate at time *t, β*_0_(*t*) is the baseline hazard function, and *γ*(*t*), *β*(*t*), *V* (*t*), and *W* (*t*) are time-varying coefficients quantifying the additive contribution of each covariate to the hazard at time *t*. This semi-parametric formulation provides a flexible alternative for time-to-event analysis by allowing covariate effects to vary over time.

Although formulated on different functions (log-survival time in the AFT framework versus instantaneous hazard function in the additive hazards model), both approaches model a survival function *S*(*t*) and enable mediation effect estimation under censored outcomes thanks to the linear representation. The Aalen specification relaxes proportional hazards assumptions by allowing time-varying additive effects, whereas the AFT model directly parameterizes survival time and avoids proportional hazards assumptions altogether. We therefore treat these two formulations as complementary modeling strategies for investigating high-dimensional mediation effects on survival outcomes.

As in the original HDMAX2 framework, statistical significance of candidate mediators in HDMAX2-surv is assessed using a max-squared test (Equation 3). Resulting *P* -values are adjusted for multiple testing using either the Bonferroni procedure to control the family-wise error rate (FWER) or false discovery rate (FDR) controlling methods such as Benjamini–Hochberg (22) or Storey’s approach (23). For selected mediators, ACMEs are then estimated based on the mediator *M* model (Equation 1) and the survival outcome *S* model (Equations 4 or 5).

## Results

### Simulations

We first compare methods on simulated high-dimensional data (Figure 1C). Our simulations generated realistic datasets with correlated DNAm blocks, binary exposure, and Cox-modeled survival outcomes (Supplementary Material 2). Consecutive CpG blocks served as causal mediators linking exposure to survival hazard. To avoid bias, survival data were simulated using Cox models that differed from the survival models implemented in HDMAX2-surv. We systematically evaluated three methods: `HDMAX2-surv.aalen`, `HDMAX2-surv.param`, and `HIMA` (9). For `HDMAX2-surv.aalen`, `HDMAX2-surv.param`, we applied a Bonferroni correction to control the FWER at 5% significance level. HIMA (9) combines genome-wide screening with per-mediator inference under a penalised Cox proportional hazards model. It performs feature selection using SIS followed by debiased LASSO and joint significance testing. Although reliance on the Cox model was not itself a selection criterion, it provides a complementary survival modeling framework relative to our AFT and Aalen specifications, allowing comparison under different modeling assumptions. Other published methods were excluded from our benchmark due to practical and methodological constraints limiting their scalability, implementation, or comparability within a high-dimensional survival mediation setting (Supplementary Material Section 1 and Table 1).

Our simulation campaign examined nine distinct analytical pipelines combining different selection (Step 1) and estimation procedures (Step 2), with systematic variation of effect sizes and sample sizes to assess method performance across realistic parameter ranges (Supplementary Material Section 2 and Supplementary Figure 1). We evaluated the performance of the method using precision scores at both Step 1 and Step 2 of the framework (Figure 1B). The results are presented for true positive count (correctly selected CpGs among the top-*m*_causal_) at Step 1 (Figure 2A) and effect prediction accuracy (statistical significance and correct effect directionality) at Step 2 (Figure 2B). Our simulation campaign revealed consistent patterns across the parameter grid. Figure 2A and Supplementary Figure 2 show precision scores in the selection stage for five effect sizes (*µ*_*α*_ and *µ*_*β*_: 0.05, 0.1, 0.5, 2, and 5), two sample sizes (150 or 300 individuals) and among 10 000 CpGs. As expected, performance of all methods increased with larger effect sizes and sample sizes. Across most scenarios, `HIMA` and `HDMAX2-surv.param` performed slightly better than `HDMAX2-surv.aalen`. At this stage, no clear advantage emerged between `HIMA` and `HDMAX2-surv.param`. We then evaluated precision scores at Step 2, which assesses whether selected CpGs both pass the significance threshold and show effect directions consistent with the simulated mediation effects (Figure 2B and Supplementary Figure 3). To perform fair and exhaustive comparison, we combined Step 2 methods with each Step 1 result, across the same parameter variations. We observed the same general trend with improved method performance as the effect size or the sample size increased. Notably, `HDMAX2-surv.param` Step 2 consistently outperformed other approaches, regardless of the Step 1 method with which it was combined. `HIMA` exhibited lower precision scores, likely due to its more conservative *P* -value outputs, which limit detection even after adjusting selection thresholds. Computational time analysis revealed that `HDMAX2-surv.aalen` was substantially slower than both `HDMAX2-surv.param` and `HIMA`, whose runtimes were roughly equivalent (Supplementary Figure 4).

**Figure 2.**
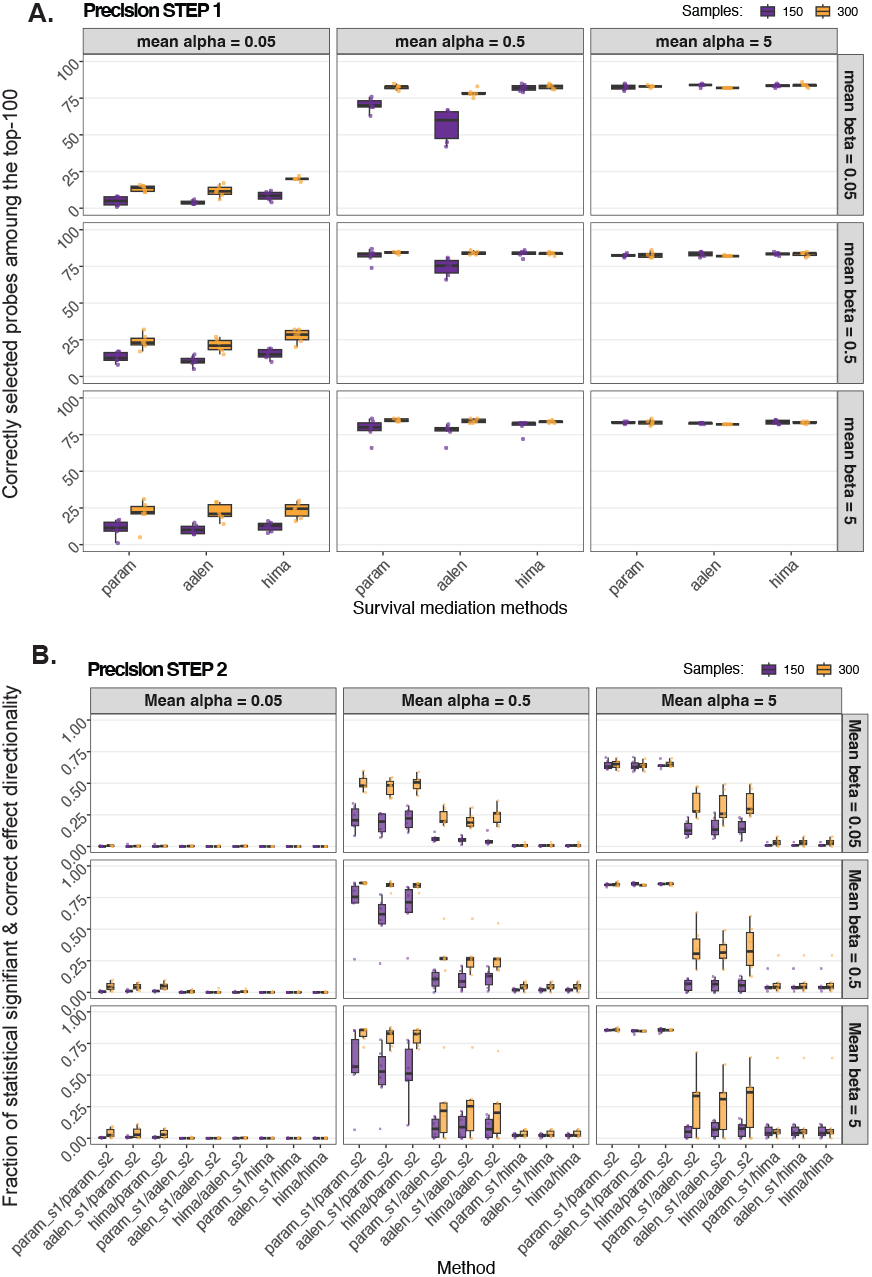
Performance evaluation of high-dimensional survival mediation methods. **(A)** Precision of methods in Step 1 (feature selection). True positive count among selected mediators, representing the number of correctly identified causal CpGs among the top-100_causal_ ranked by |*αβ*| for `HIMA` and max{*P*_*α*_, *P*_*β*_}^2^ for HDMAX2-surv. **(B)** Precision of methods in Step 2 (mediation analysis). Proportion of CpGs selected in Step 1 whose mediation effects in Step 2 exhibit both statistical significance and correct effect directionality. For both **(A)** and **(B)**, simulation parameters: *m*_causal_ = 100 causal CpGs among *p* = 10 000 total CpGs. *X* → *M* effects: *µ*_*α*_ ∈ {0.05, 0.5, 5*}, σ*_*α*_ = 0.1. *M* → *S* effects: *µ*_*β*_ ∈ {0.05, 0.5, 5*}, σ*_*β*_ = 0.1. Overlap proportion: *π*_overlap_ = 0.8. Two sample sizes are shown (*n* = 150, purple; *n* = 300, orange) across 6 replications.

As a control, we examined the behavior of each method in detecting false positives by running simulations with either *α* = 0 (Supplementary Figure 5) or *β* = 0 (Supplementary Figure 6). When *β* = 0, `HIMA` displayed a high false positive rate, while smaller numbers of false positives were observed in `HDMAX2-surv` methods. These false positive rates are partially explained by the feature selection approach, which ranks potential mediators using scores that combine both pathway effects: |*α*_*j*_*β*_*j*_ | for `HIMA` and max{*P*_*α*_, *P*_*β*_*}*^2^for `HDMAX2-surv`. Consequently, a strong effect on one pathway (*α* or *β*) can drive the ranking even when the other pathway has no effect (Supplementary Figure 7). Our observations indicate that HIMA is more susceptible to this issue than HDMAX2-surv, likely due to its multiplicative scoring approach that amplifies single-pathway effects.

Overall, `HDMAX2-surv.param` delivered the best performance: it matched `HIMA` in Step 1 selection and computational efficiency while providing substantially higher accuracy in effect estimation and lower false positive rates. Based on these simulation results, we selected `HDMAX2-surv.param` for downstream analysis of real data.

### Mediation of smoking status on PDAC survival

Epigenetic modifications, such as DNA methylation (DNAm), are increasingly recognized as key mediators of environmental exposures on disease progression. In cancer, DNAm patterns reflect both exposure history and tumor biology, influencing clinical outcomes through complex pathways (24). For instance, tobacco exposure alters genome-wide DNAm patterns which may in turn affect survival via immune microenvironment remodeling (18; 25). Although the relationship between tobacco exposure and PDAC incidence has been extensively characterized, the causal mechanisms linking smoking with the survival outcome in PDAC patients remain poorly understood. To address this knowledge gap, we investigated the effect of smoking on survival among PDAC patients (conditional on disease status), specifically examining how DNAm patterns mediate this effect. We used the comprehensive multi-omic dataset from The Cancer Genome Atlas (TCGA; 26), which provides integrated genomic, transcriptomic, and epigenetic profiles for PDAC patients along with detailed clinical characteristics, including smoking status, age, sex, tumor grade, and survival outcome (Supplementary Table 3 and Supplementary Figure 8). Our analysis focused on a subset of 112 patients with confirmed PDAC diagnosis, as previously established in (27). Detailed description of clinical variables and inclusion criteria are provided in the Supplementary Material (Section 3.1). We first evaluated the association between tobacco exposure and overall survival using a Cox proportional hazards model adjusted for age, sex, and tumor grade (*n* = 112; 59 events). After adjustment, the total effect of tobacco exposure was not significantly associated with survival (HR = 0.93, 95% CI [0.51-1.68], *p* = 0.81). In mediation analysis however, the absence of a statistically significant total effect does not preclude the presence of indirect effects, supporting the hypothesis that smoking effect on prognosis may operate through complex intermediate molecular mechanisms (28).

We then applied the `HDMAX2-surv.param` method to characterize the causal relationships between tobacco exposure, DNAm, and patient survival (Supplementary Material Section 3.2 and Supplementary Figure 9). A total of 371,033 CpG sites were considered as possible mediators. Using FDR control at 10% threshold, we identified only 10 mediating CpG sites (Figure 3A-top and Supplementary Figure 10), illustrating our statistical power limitations. To address this, we adopted a spatial aggregation strategy using `comb-p` (29) to combine adjacent *P* - max^2^ values from individual CpGs identified in Step 1 of `HDMAX2-surv.param`, then selected the top 50 candidate regions (Figure 3A-bottom). Because CpGs co-localized within 1 kb typically exhibit strong local correlation and shared regulatory context, we summarized Aggregated Mediator Regions (AMRs) using the mean DNAm across constituent CpGs. Given the strong local correlation structure of methylation within short genomic distances, regional averaging provides a robust and biologically coherent summary while reducing probe-level noise and dimensionality; in cases of discordant CpGs, averaging acts conservatively by attenuating opposing effects. From the top 50 candidate regions, we identified 36 AMRs with statistically significant Average Causal Mediation Effects (ACMEs; Figure 3B). The identified AMRs exhibited heterogeneous effects on patient survival. We identified 5 deleterious AMRs and 31 protective AMRs. Detailed characteristics of AMRs with significant ACMEs are presented in Supplementary Table 4. Since a specific parametric distribution must be specified in the AFT model for each AMR, ACME values are not strictly comparable across AMRs when different distributional assumptions (Weibull, log-normal, log-logistic) are applied. Notably, some AMRs show substantial variability in effect size estimates, spanning several orders of magnitude with correspondingly wide confidence intervals. For example, AMR50 yields an ACME of -123.5 [95% CI: -285.1, - 6.4]. Such wide intervals may reflect convergence issues in the bootstrap procedure and the limited sample size, underscoring the preliminary nature of these estimates. This variability highlights the exploratory scope of our analysis and the critical importance of experimental validation. While the direction of effects may be more reliable than their precise magnitudes, future studies with larger sample sizes will be required to refine these estimates and reduce uncertainty. Among the significant AMRs, we identified four genes previously associated with PDAC occurrence or outcome, defined as having more than two prior citations in PubMed (located in AMR2, 5, 17 and 37; see Supplementary Material section 3.2 and Supplementary Figure 11). These four AMRs demonstrated distinct mechanistic mediation patterns, yet overall showed a protective effect on survival. However, not all effects were protective. In some cases, antagonistic patterns resulted in deleterious outcomes, such as AMR38, where tobacco-induced hypomethylation shifts the survival curve backward and reduced survival, or AMR25, where tobacco-induced hypermethylation had also a deleterious impact. These findings reveal complex, context-dependent relationships between tobacco exposure and survival outcomes, suggesting that DNA methylation alterations may both amplify and attenuate tobacco-related survival effects depending on the specific genomic locus. Beyond the established candidates we identified, the biological mechanisms underlying the remaining AMRs represent promising avenues for future mechanistic investigation.

**Figure 3.**
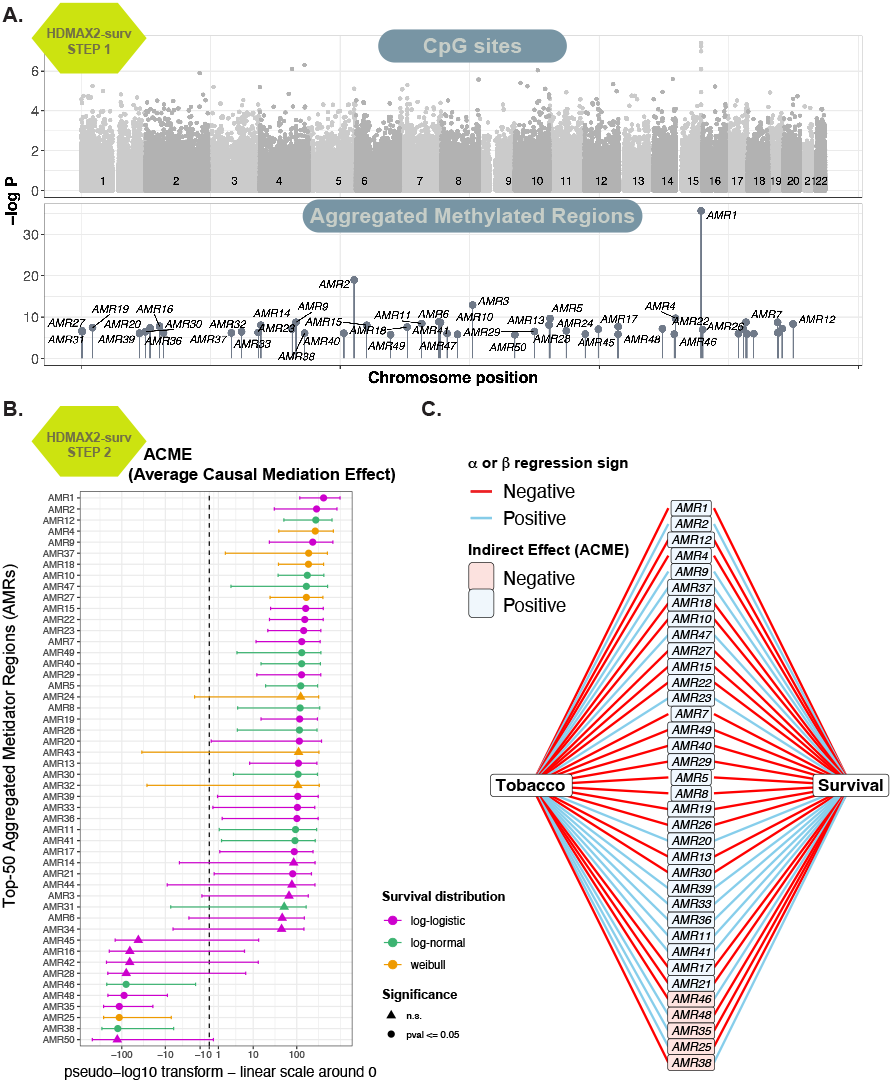
Mediation analysis of the effect of smoking (tobacco exposure *T*) on survival *S* of PDAC patients through DNAm *M*. **(A)** Top: Manhattan plot of DNAm CpG − log_10_(*p*) values from HDMAX2. Bottom: Manhattan plot of − log_10_(*p*) values for top-50 candidate Aggregated Methylated Regions (AMR). **(B)** Indirect effects (ACME) for each AMR; point= estimate, bar= 90 %CI. Data were transformed using the `transform_pseudo_log` of the ‘scales’ R package (base = 10, sigma = 1), switching to a linear scale for absolute values below 1. Colors indicate the survival distribution retained in the AFT models, for each AMR. **(C)** Decomposition of significant ACME of smoking on survival: 31 AMRs show a positive indirect effect and 5 a negative effect. Boxes colours reflect the sign of the ACME (blue = protective, red = deleterious) and segment colours indicates the direction of the effects *α* and *β*, for each model *T* → *M* and *M* → *S*.

### Causal discovery of tobacco-survival mechanisms

We leveraged HDMAX2-surv to dissect multi-layered mechanisms linking tobacco exposure to PDAC prognosis, focusing on complex interactions among tumor immune infiltration, DNAm, and patient survival, a relationship supported by prior evidences. First, immune cell composition has been consistently linked to survival outcomes in PDAC (25; 30). Second, Weissman *et al* (18) suggest that immune infiltration within the tumor microenvironment mediates the effects of tobacco exposure on survival in PDAC patients. To effectively identify and interpret causal mechanisms while integrating biological prior knowledge, we implemented an integrated analytical pipeline combining immune cell deconvolution, causal discovery, and serial mediation analyses (Figure 4). This demonstrates the power and versatility of the HDMAX2-surv framework.

**Figure 4.**
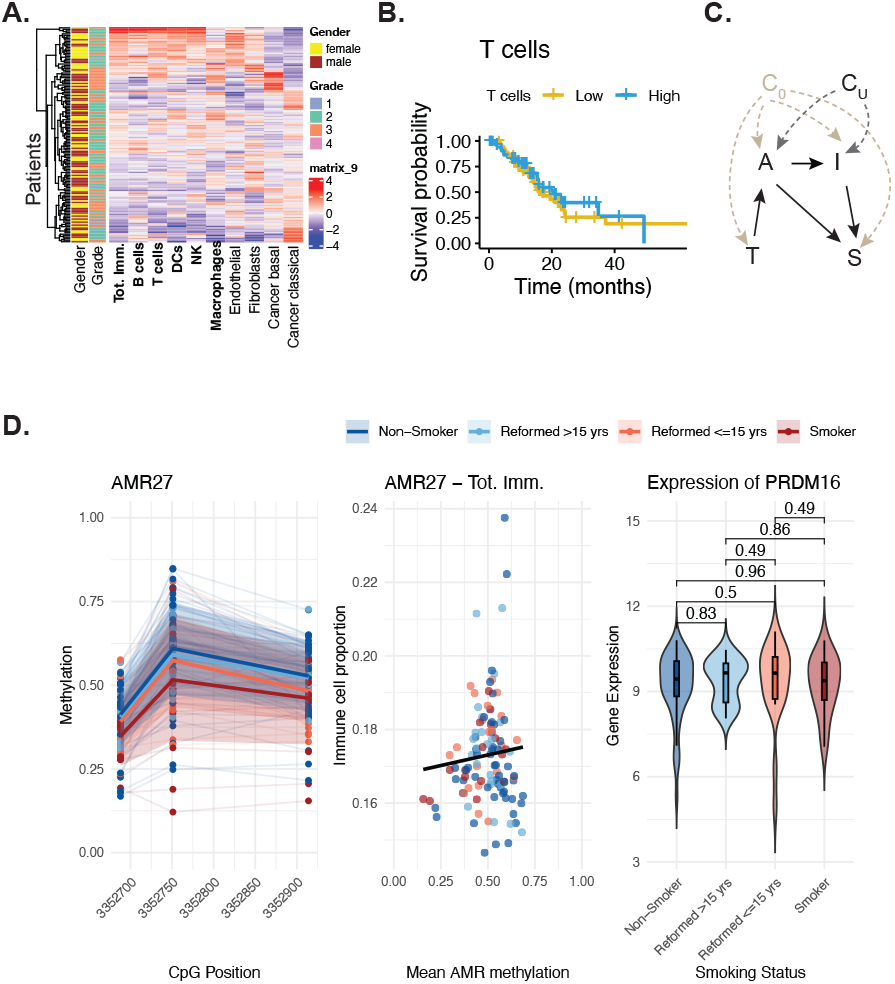
Integrated analysis of immune-mediated epigenetic pathways in PDAC survival. **(A)** Standardized deconvolution estimates for 9 cell types in TCGA-PAAD (n=112). Rows represent samples clustered by immune composition. *Tot.Imm*. indicates total immune infiltration. **(B)** Kaplan-Meier survival curves for T cell infiltration (high vs. low, median split). Blue: high infiltration; orange: low infiltration. **(C)** Directed acyclic graph (DAG) framework showing retained (green) and removed (red) relationships after unconditional independence testing. *C*_*O*_: observed confounders; *C*_*U*_: latent factors. **(D)** Example methylation patterns for *PRDM16* -AMR27: (i) CpG-specific methylation by smoking status, (ii) correlation with T cell infiltration, (iii) gene expression by smoking category. Colors: red=smokers, blue=non-smokers.

We first quantified immune infiltration (Figure 4A) using a consensus approach integrating four deconvolution methods: RLR (31), SCDC (32), InstaPrism (33), and MuSiC (34) (Supplementary Figure 12 and Supplementary material 4.1). Then we developed a causal discovery framework to investigate immune-mediated pathways linking tobacco exposure, DNAm, and survival in PDAC patients. Starting from a fully connected directed acyclic graph (DAG) including tobacco exposure (*T*), methylation levels of significant AMRs (*A*), immune infiltration components (*I*), and survival (*S*) (Supplementary Material section 4.2 and Supplementary Figure 13), we removed possible causal links by sequentially testing unconditional and conditional independence among variables while adjusting for observed confounders (age, sex, tumor grade) and latent factors derived from HDMAX2-surv (Supplementary Figure 14). Precisely, Cox proportional hazards modeling revealed significant associations between some immune cell types and survival (Figure 4B; Supplementary Figure 15). For example, T cell infiltration showed a trend toward improved survival (HR=0.58, 95% CI[0.28–1.19], p=0.140). Causal direction were first oriented using biological prior knowledge: *T* → *A, A* → *S, I* → *S*. Then AMR-immune pairs were systematically assessed via statistical models to infer causal directions. Conditional independence tests determined whether AMRs influenced survival directly or solely via immune infiltration, leading to two candidate DAGs. Logistic modeling further clarified the directionality between AMRs and immune variables (*A* → *I*) by testing conditional independence of *I* from *T* given *A* (Figure 4C). In total, 29 AMR-immune pairs supported direct effects on both immune infiltration and survival, while 2 pairs acted only through immune-mediated pathways, highlighting the complex, multi-layered causal mechanisms of tobacco on PDAC prognosis (Supplementary Figure 16).

Next, we sought to identify whether indirect effects exist between tobacco exposure and PDAC patient survival through identified mediator chains (*A* → *I* → *S*). Using a counterfactual approach adapted from Zugna et al. for parametric survival models (35), we analyzed the 29 AMR-immune pairs identified by causal discovery (Supplementary Material 4.3 and Supplementary Figure 17). Among these, seven AMRs exhibited statistically significant total indirect effects mediated via immune infiltration (AMR4, AMR5, AMR9, AMR17, AMR20, AMR23, and AMR27, described in Supplementary Figure 18, and 19). Notably, several AMRs revealed a paradoxical protective effect of tobacco exposure on survival, most prominently AMR27. This region is located in the gene body of PRDM16, a zinc finger transcription factor known for recurrent translocations in acute myeloid leukemia (36) but not previously linked to PDAC. This locus is normally hemi-methylated but shows a smoking-associated demethylation bias (Figure 4D). Notably, smoking status was not associated with changes in PRDM16 expression (Wilcoxon rank-sum test; Figure 4C), suggesting an epigenetic-specific mechanism that might be acting on other loci. Across all selected AMRS, we observed complex effects of immune infiltration, consistent with prior literature. For instance, chronic pancreatitis influences PDAC outcomes, probably through remodeling of the immune microenvironment, but the mechanisms remain complex and not fully understood (37). Such scenarios highlight the importance of mediation analysis in dissecting underlying biological mechanisms even when overall associations appear non-significant.

Our results demonstrate how the HDMAX2-surv framework enables scalable, integrated analyses in cancer epigenomics. In PDAC, tobacco exposure may exert dual effects on survival: protective via specific epigenetic-immune pathways (e.g., AMR27), or deleterious through alternative mechanisms. This complex interplay underscores the need for integrated analytical approaches to disentangle multi-layered mediation mechanisms in cancer prognosis.

## Conclusion

In this study, we developed a novel high-dimensional mediation analysis method for survival outcomes and applied it to investigate the role of DNA methylation in mediating tobacco effects on PDAC patient survival.

Our HDMAX2-surv method addresses a critical gap in high-dimensional mediation analysis for survival data. Through extensive simulations, we demonstrated superior performance compared to existing approaches, particularly in handling the scale and complexity of epigenome-wide DNAm data. The parametric survival modeling approach offers computational advantages while maintaining robust statistical inference, although it remains dependent on the correct model specification. Additional simulations exploring diverse effect size distributions, inter-probe correlations, and DNAm noise structures would further strengthen its evaluation.

Our findings also show that current methods claiming to handle high-dimensional mediation may not remain robust in genuinely large-scale settings, highlighting the need for methodological advances. This is particularly important for DNAm mediation analysis, which is increasingly applied in studies of chronic disease and environmental exposures (38). Beyond DNAm, the proposed framework can be adapted to other exposure–outcome contexts where epigenetic mechanisms are implicated. Estimating total mediated effects in survival analysis remains a complex challenge, particularly when mediators exert opposing influences, as illustrated in our tobacco–survival analysis. Developing robust strategies for such settings should be a priority for future methodological research.

We also developed an ad hoc framework to identify causal relationships and perform serial mediation analysis between AMRs and immune infiltration. Several limitations of this approach warrant consideration. First, accurate estimation of direct and indirect effects in serial mediation relies on the assumption of no unmeasured confounders across the exposure–mediator–outcome relationships (35). Although our causal discovery framework incorporated measured confounders (age, sex, tumor grade) and verified the independence of latent factors from tobacco exposure, the potential influence of unmeasured confounders—such as genetic predisposition, environmental exposures, or lifestyle factors—remains a limitation that could bias our causal effect estimates. Second, our framework assumes linear relationships between intermediary molecular variables, which may not capture the full complexity of biological interactions (39).

Given the modest cohort size, statistical power and false discovery control were major challenges. Our multi-stage analytical pipeline, spanning CpG screening, AMR construction, mediation testing, causal discovery, and serial mediation, may increase the risk of error propagation. Rather than aiming for exhaustiveness, we adopted a candidate-driven strategy designed to prioritize biologically plausible regions for downstream validation. Although significance thresholds inevitably retain some arbitrariness, the consistency and biological coherence of identified candidates provide support for the validity of our findings.

Mediation analysis, while classically used in epidemiology, remains underutilized in oncology research. PDAC is known to be refractory to immunotherapy, but the underlying mechanisms have yet to be fully characterized (40). Our work opens new perspectives on the mechanisms underlying tobacco effects on survival, highlighting specific AMR-immune cell combinations that warrant experimental validation. The identified loci may represent promising candidates for diagnostic and prognostic evaluation, pending functional validation. Furthermore, our findings suggest that epigenetic-immune interactions may serve as actionable biomarkers for personalized treatment strategies in tobacco-exposed PDAC patients.

In conclusion, this study presents HDMAX2-surv, a new method to perform high-dimension mediation analysis of survival data. It demonstrates the power of integrating high-dimensional mediation analysis with causal discovery approaches to uncover complex biological mechanisms. Applied to PDAC, our approach reveals previously uncharacterized epigenetic–immune interactions potentially linking tobacco exposure to patient prognosis, offering both methodological and biological insights for future cancer research.

## Data availability

The code for data analysis and figure generation is available on GitHub (https://github.com/bcm-uga/tims-pdac).

## Author contributions statement

Conceptualization and methodology: F.P., E.A., E.D, A.S, N.V. and M.R., Project administration: M.R. Data analysis: F.P., E.A, N.V. and M.R., Writing: F.P., E.A. and M.R. All authors have reviewed the manuscript and agreed to its publication.

## Acknowledgments

We thank Yuna Blum for critical reading of the manuscript. We thank the GRICAD UGA mesocenter for computing resources. We thank Hugo Barbot and Olivier François for insightful scientific discussions.

## References

1. K. Imai, et al., Psychological Methods (2010).

2. D. Tingley, et al., Journal of Statistical Software (2014).

3. M. G. B. Blum, et al., Environmental Health Perspectives (2020).

4. L. Lapointe-Shaw, methodology (2018). et al., BMC medical research

5. S.-H. Lin, et al., Statistics in medicine (2017).

6. M. E. Sobel, Sociological Methodology (1982).

7. R. M. Baron, et al., Journal of Personality and Social Psychology (1986).

8. C. Luo, et al., PLOS Computational Biology (2020).

9. H. Zhang, et al., Bioinformatics (Oxford, England) (2021).

10. S. Chi, et al., The Annals of Applied Statistics (2024).

11. Z. Shao, et al., PLOS Computational Biology (2021).

12. P. Tian, et al., Bioinformatics (2022).

13. L. Huang, et al., Bioinformatics (2023).

14. S. Ahn, et al., ArXiv (2025).

15. F. Pittion, et al., Peer Community in Genomics (2025).

16. B. Jumentier, et al., Environmental Health Perspectives.

17. E. Afghani, et al., Hematology/Oncology Clinics of North America (2022).

18. S. Weissman, et al., Pancreas (2020).

19. S. Iodice, et al., Langenbeck’s Archives of Surgery (2008).

20. S. M. Lynch, et al., American Journal of Epidemiology (2009).

21. N. Momi, et al., Carcinogenesis (2012).

22. Y. Benjamini, et al., Journal of the Royal Statistical Society. Series B (Methodological) (1995).

23. J. D. Storey, Journal of the Royal Statistical Society Series B: Statistical Methodology (2002).

24. S. Rousseaux, et al., BMC Medicine (2020).

25. S. Sivakumar, et al., Nature Communications (2025).

26. J. N. Weinstein, et al., Nature Genetics (2013).

27. R. Nicolle, et al., Cancers (2019).

28. H. P. O’Rourke, et al., Journal of Studies on Alcohol and Drugs (2018).

29. B. S. Pedersen, et al., Bioinformatics (2012).

30. P. Gomez-Rubio, et al., Gut (2017).

31. S. C. Zheng, et al., Nature Methods (2018).

32. M. Dong, et al., Briefings in Bioinformatics (2021).

33. M. Hu, et al., Bioinformatics (2024).

34. X. Wang, et al., Nature Communications (2019).

35. D. Zugna, et al., BMC Medical Research Methodology (2022).

36. I. Nishikata, et al., Blood (2003).

37. K. K. Mahadevan, et al., Science (2024).

38. R. Fujii, et al., Epigenetics (2022).

39. Q. Cai, et al., Genome Research (2024).

40. F. Cappellesso, et al., Nature Cancer (2022).

