## Supplementary Material for "HDMAX2-surv: high-dimensional mediation analysis of survival data with application to pancreatic cancer"

March 4, 2026

### 1 Method

#### 1.1 Description of published methods dedicated to survival mediation analysis

| Method | Survival model | STEP1 (Feature selection) | STEP2 (Mediation analysis) | Reason for inclusion / exclusion in our benchmark |
|---|---|---|---|---|
| **hima-survival**<br>Zheng et al., 2021 | Cox | ✓<br>SIS | ✓<br>Joint test | **Included** for STEP1 and STEP2 comparisons. |
| **mediateR**<br>Huang et al., 2023 | AFT (survreg) | ×<br>—<br>(no built-in screening) | (✓)<br>Ridge-penalised AFT | Requires $\geq 3$ exposures and a causal-path matrix; failed on $p = 10\,000$ probes: **excluded**. |
| **SMAHP**<br>Ahn et al., 2025 | AFT | (✓)<br>MCP-penalised AFT, then SIS | (✓)<br>Joint significance test | Designed for a minimum of $n = 3$ multiple exposures: **excluded**. |
| **CoxMKF**<br>Tian et al., 2022 | Cox | ✓<br>Aggregated model-X knockoff filter | ×<br>— | No indirect-effect estimation framework ; Implementation errors prevented application: **excluded**. |
| **multimediate**<br>Domingo-Relloso et al., 2024 | Additive Aalen | ×<br>— | ✓<br>Additive Aalen mediation + quasi-Bayesian ACME IC | No high-dimensional selection; used only for effect estimation in our workflow—thus not relevant for selection comparison: **excluded**, but the HDMAX2_SURV_AALEN method derives from the multimediate method. |
| **IUSMMT**<br>Shao et al., 2021 | Cox | ×<br>Single mediator | (✓)<br>intersection union mixture test | Designed for multiple DNA-methylation exposures and a single mediator; not applicable to our single-exposure, high-dimensional mediator setting: **excluded**. |
| **MASH**<br>Chi et al., 2024 | Cox | ?<br>SIS + MCP + BH | ?<br>$R^2$-based mediation measure | No publicly available R / Python implementation; not reproducible in our pipeline: **excluded**. |

**Supplementary Table 1: Candidate mediation methods with survival outcomes considered, and justification for their inclusion or exclusion in the simulation study. STEP1 indicates availability of high-dimensional feature selection; STEP2 indicates availability of mediation effect estimation. ✓= fully available; (✓) = available with limitations; ×= not available; ? = unknown due to implementation unavailability.**

#### 1.2 Rationale for method exclusion

Several alternative high-dimensional mediation methods were not retained for benchmarking due to practical or methodological constraints. Although mediateR (1) and SMAHP (2) are presented as high-dimensional approaches, they have primarily been evaluated on moderate-sized datasets and lack scalable variable selection procedures suitable for genome-wide CpG analyses. CoxMKF (3) is restricted to knockoff-based selection and does not provide mediation effect estimates, precluding direct comparison. MASH does not offer publicly available `R` or `Python` implementations. IUSMMT is designed for multiple DNA methylation exposures with a single mediator, and multimediate (survival) (4) requires a pre-specified set of mediators, preventing genome-wide screening. Although some of these methods could theoretically be incorporated into the effect estimation stage of our workflow, their current implementations do not allow systematic side-by-side benchmarking (Supplementary Table 1).

### 2 Simulation framework

To evaluate HDMAX2-surv performances and compare them with state-of-the-art method, we performed simulations. We simulated datasets mirroring our mediation setting with DNAm and a survival outcome.

#### 2.1 Objectives and design

The goal of this simulation study is to assess high-dimensional mediation methods with a survival outcome. For each method, we evaluate:

- its ability to select mediators (ranking and selection thresholds);
- its ability to estimate effect sizes.

We simulate the following variables:

- the exposure $X$ (continuous, with an optional dichotomized version $X_{\text{bin}} = \mathbf{1}_{\{X \geq 0\}}$ used in some scenarios);
- the methylation matrix $M$ incorporating
  - the influence of latent factors $U$,
  - correlation with the exposure through the $X \to M$ pathway,
  - within-block correlation among probes,
  - Gaussian noise;
- The **survival time** $T$ is drawn from an exponential model, with a hazard depending on both methylation and exposure ($M \to T$, $X \to T$).
- The **censoring time** $C$ is sampled independently from a Gamma distribution and calibrated to yield approximately 20% censoring.

We generate complete datasets $(X, M, T, C)$ for $N$ individuals and $p$ CpG sites arranged into 16 correlated blocks (the within-block correlation is fixed at 0.5). Each individual has $K$ unobserved latent factors $\mathbf{U}_i = (U_{i1}, \ldots, U_{iK})$.

**1. Causal blocks and effects $\alpha$ and $\beta$.** We randomly select $b$ blocks of 10 probes. For each selected block, the exposure-to-mediator effects and mediator-to-outcome effects are drawn as

$$\alpha_j \sim \mathcal{N}(\mu_\alpha, \sigma_\alpha^2), \qquad \beta_j \sim \mathcal{N}(\mu_\beta, \sigma_\beta^2),$$

with a *block-level sign* $s \in \{-1, 1\}$ sampled once per block and applied to both $\alpha_j$ and $\beta_j$ to mimic hypo- vs. hyper-methylated regions. The intersection of indices where $\alpha_j \neq 0$ and $\beta_j \neq 0$ (proportion $\pi_{\text{overlap}}$) defines the set of *true mediators*. The overlap proportion $\pi_{\text{overlap}}$ controlled the degree to which the sets of CpGs with non-zero coefficients $\alpha_j$ and non-zero $\beta_j$ coincided. Specifically, $(1-\pi_{\text{overlap}})/\pi_{\text{overlap}} \times m_{\text{causal}}$ CpGs had non-zero exposure effects ($\alpha_j \neq 0$) but no survival effects ($\beta_j = 0$), and vice versa. The intersection of these two sets defined the *true mediators*, carrying both exposure-to-mediator and mediator-to-outcome effects.

**2. Exposure and latent factors.**

We jointly simulate $(\mathbf{U}_i, X_i)$ from a $(K+1)$-variate normal distribution, and tune the loadings so that a pre-specified variance-share parameter $\nu_X \in (0,1)$ controls how strongly the latent confounders $U$ explain the variability of the exposure $X$. Specifically, $\nu_X$ represents the proportion of variance in $X$ attributable to its linear dependence on $U$, while the direction of this dependence is randomly drawn at each replicate to reflect stochastic variability.

In some scenarios we dichotomize the exposure as $X_{\text{bin}} = \mathbf{1}_{\{X \geq 0\}}$.

**3. Methylation matrix.** For each site $j$ and individual $i$,

$$M_{ij} \;=\; \mathbf{U}_i \mathbf{V}_j^\top \;+\; X_i A_j \;+\; \varepsilon_{ij}, \qquad \varepsilon_{ij} \sim \mathcal{N}(0, \sigma^2), \quad \mathbf{V}_j \sim \mathcal{N}(\mathbf{0}, \sigma_V^2 I_K).$$

The 16 correlated blocks among probes are imposed via a block-structured correlation matrix, after which the values are mapped to the unit interval $(0,1)$ to obtain beta-like methylation levels.

**4. Survival time and censoring.** Survival times follow an exponential model with baseline hazard $\lambda_0$ and linear predictor combining mediator and exposure effects:

$$T_i \sim \text{Exp}\Big(\lambda_0 \exp\{\mathbf{M}_i^\top \mathbf{B} \;+\; \theta\, X_i\}\Big),$$

where $\mathbf{B} = (B_1, \ldots, B_p)$ and $\theta$ denotes the *direct effect of $X$ on $S$* (log-hazard ratio).

Independent censoring times $C_i \sim \text{Gamma}(k = 2, \text{scale})$ are calibrated to yield approximately 20% censoring. We observe $O_i = \min(T_i, C_i)$ and the event indicator $\delta_i = \mathbf{1}_{\{T_i \leq C_i\}}$.

#### 2.2 Choice and rationale for parameters

Here we give a comprehensive description of the parameters we wished to explore. The parameter grid retained for the manuscript is narrower; we progressively refined it across versions and interim results. Sample sizes were chosen to match our data ($n = 150$) and to include a more statistically powered scenario ($n = 300$).

The number of potential mediators was set to 1,000 in pilot runs and 10,000 for the main comparisons.

Mean effect sizes were initially chosen somewhat arbitrarily but within a scale consistent with our empirical data. Effects $A$ (exposure $\rightarrow$ CpG) are more directly interpretable, whereas for $B$ (CpG $\rightarrow$ survival) the impact is less direct because it enters the survival-time generation model (Section 2.1). We also allowed the standard deviations of the effects to vary, following the same logic as for the means.

We examined several levels of overlap among causal probes, from 50% to 100%.

Finally, the compared methods are not parameters in the strict sense—they are the objects of comparison—but including them in the table helps clarify the combinations explored.

For each retained combination we generate between 5 and 10 replicates. Each replicate is produced with a distinct pseudo-random seed (seed = 1, 2, ..., $R$), ensuring reproducibility while producing distinct datasets.

| Parameter | Levels examined |
|---|---|
| Total number of CpG $p$ | 1 000; 10 000 |
| Sample size $N$ | 150; 300 |
| Mean effect exposure $\to$ CpG ($\mu_\alpha$) | 0.05; 0.10; 0.50; 2; 5 |
| Mean effect CpG $\to$ survival ($\mu_\beta$) | 0.05; 0.10; 0.50; 2; 5 |
| Effect variance ($\sigma_A^2 = \sigma_B^2$) | 0.01; 0.05; 0.10 |
| Overlap $A \cap B$ ($\pi_{\text{overlap}}$) | 0.50; 0.80; 1 |
| Number of causal CpGs ($m_{\text{causal}}$) | 40; 80; 100 |
| Number of causal blocks | 4; 8; 10 |
| Number of latent factors $K$ | 5 |
| Proportion of variance in $X$ attributable to its linear dependence on $U$ (used to calculate $\nu_X$) | 0.2 |
| $U$ standard deviation (used to calculate $\nu_X$) and $V$ standard deviation | 0.1 |
| Methods compared | `HDMAX2_param`; `HDMAX2_aalen`; `hima` |

**Supplementary Table 2: Full grid of parameters explored in the simulation study.**

#### 2.3 Simulation campaign

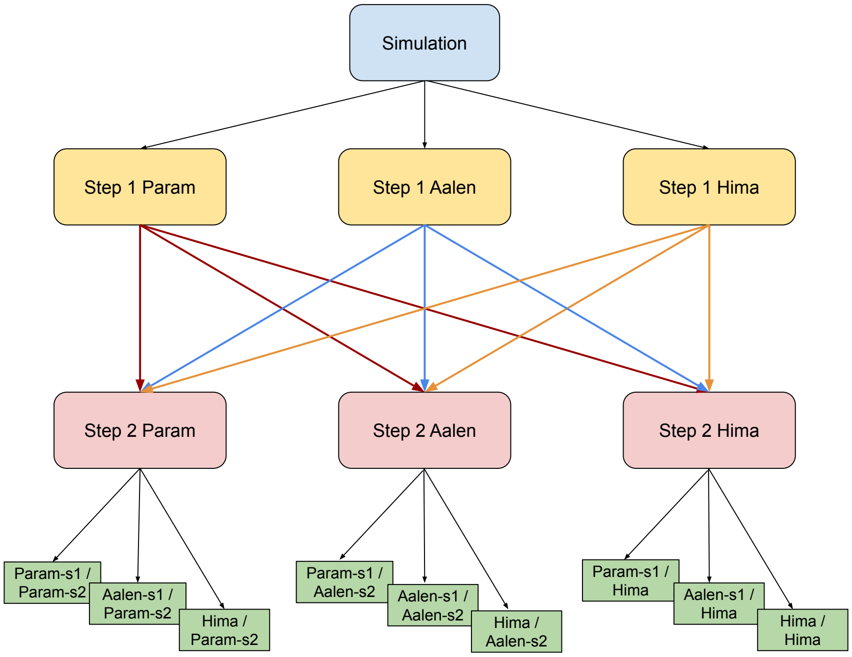


**Supplementary Figure 1: Simulation workflow DAG**

We explored a range of simulation settings defined by varying the mean exposure $\to$ methylation effect ($\mu_\alpha$), the mean methylation $\to$ survival effect ($\mu_\beta$), and the sample size ($n$). For each parameter combination, we ran the selection stage **step 1** (figure 1B) using three procedures, `run_AS_surv_param()` (parametric survival), `run_AS_surv_aalen()` (Aalen additive), and `HIMA_survival()`. We retained the 100 CpGs with the smallest $P$-values computed by the HDMAX2 max-squared test (5) for the two first procedures and set the HIMA procedure to select 100 CpGs. In the estimation stage **step 2** ((figure 1B)), we quantified indirect effects for the top-$m_{\text{causal}}$ candidates ($m_{\text{causal}}$ equal to the true number of simulated causal CpGs): `estimate_effect_surv_param()` (using the R package `mediation` (6)) for the parametric branch and `estimate_effect_surv_aalen()` (using the R package `multimediate_survival` (4), (7)) for the Aalen branch. HIMA integrates selection and estimation in a single call. We adjusted its FDR threshold to return exactly $N$ probes and treated its output as directly comparable **step 2** results. Combining the three selectors with the three estimators gave nine distinct **step 2** result sets, summarised in and in the workflow Directed Acyclic Graph (see Supplementary Figure 1).

#### 2.4 Performances evaluation

We evaluated the performance of Step 1 (mediator selection) using the "precision score" defined as the count of True Positives (TP) among selected mediators, representing the number of correctly identified causal CpGs among the top-$m_{\text{causal}}$ ranked candidates, where $m_{\text{causal}}$ corresponds to the number of causal simulated CpGs. This metric directly assesses the method's ability to prioritize true mediators over non-mediators in high-dimensional settings. We assessed Step 2 performance using a "precision score" that requires both statistical significance and correct effect directionality. Specifically, a true prediction was validated when: (i) the Average Causal Mediation Effect (ACME) was correctly identified as statistically significant and (ii) the estimated effect sign matched the simulated ground truth. Among the top 100 CpGs selected in Step 1, we retained those with significant estimated effects using a Bonferroni-corrected threshold ($\alpha = 0.05/100 = 0.0005$) and verified whether their estimated effects were consistent with the simulated mediation effects. This stringent evaluation ensures that the methods not only detect mediation, but also provide accurate effect estimates.

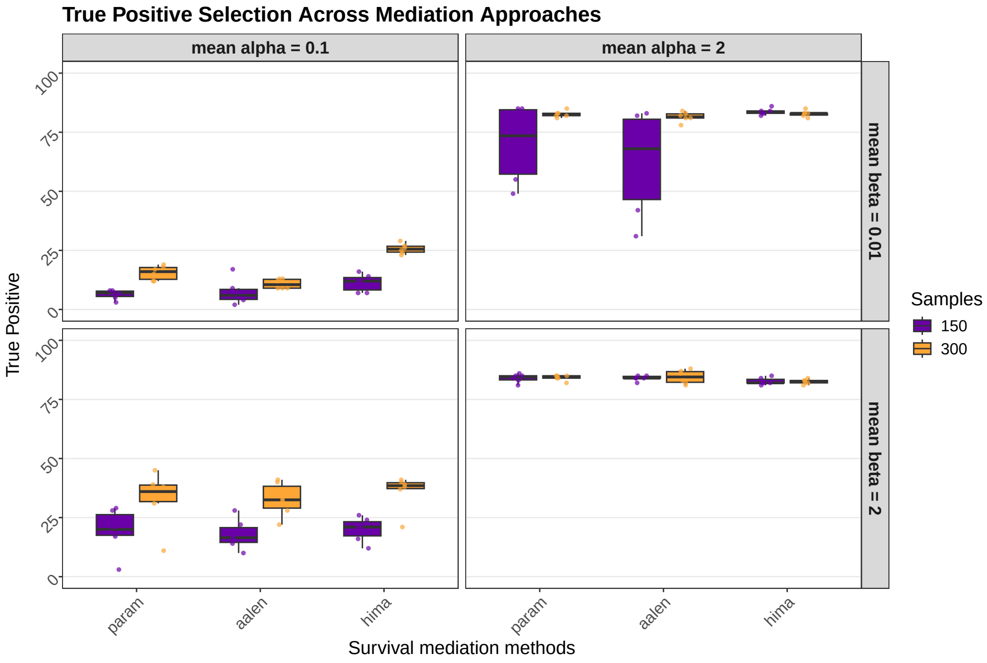


**Supplementary Figure 2: Distribution of True Positive count (across 5 replications) for two mean $\alpha$ effect sizes (0.1, 2) and two mean $\beta$ sizes (0.01, 2) and two sample sizes (n = 150, purple; n = 300, orange)**

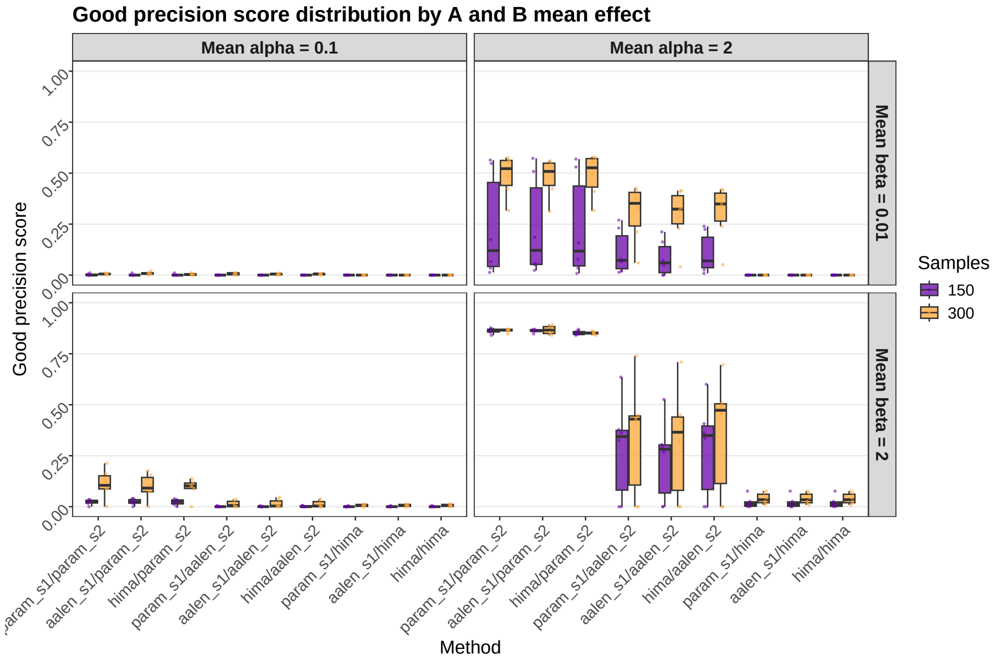


**Supplementary Figure 3: Good-precision score—the proportion of probes whose mediation effect in step 2 is both significant and has the correct sign, presented for the same simulation settings. Distribution of F1-scores (across 5 replications) for two mean $\alpha$ effect sizes (0.1, 2) and two mean $\beta$ sizes (0.01, 2) and two sample sizes (n = 150, purple; n = 300, orange)**

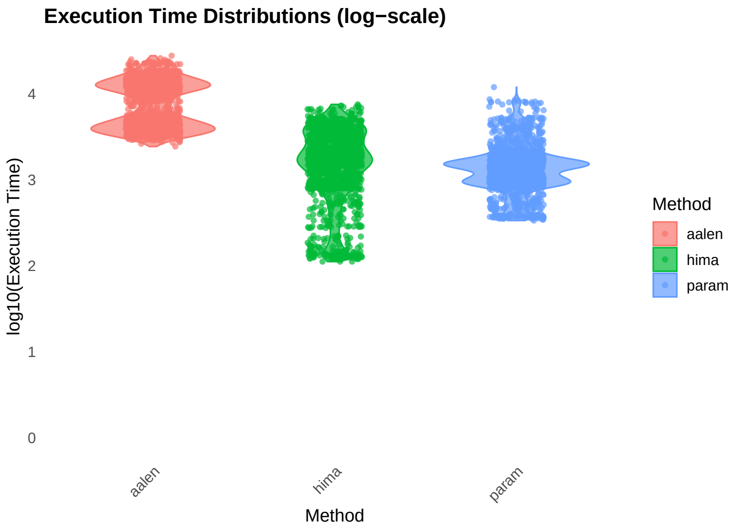


**Supplementary Figure 4: Computation time of each method. Log-scale computation times for each method, pooled over all scenarios.**

#### 2.5 Behaviour under the null hypothesis

To compare the behaviour of each survival model when mediation analysis is conducted on a high-dimension data with no true mediators, we evaluated the step 1 on simulations with effect parameters set to 0 for exposure → mediator effect or mediator → outcome effect (see Supplementary Figures 5, 6 and 7).

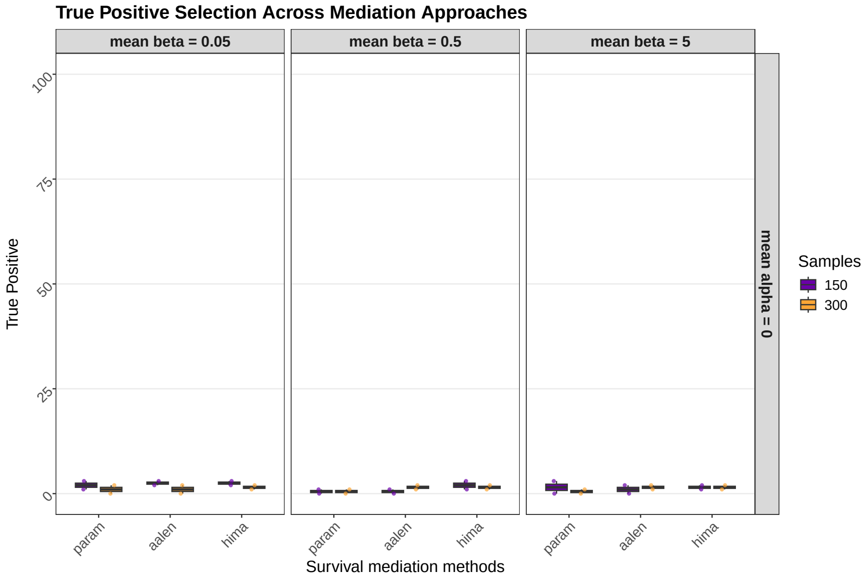


**Supplementary Figure 5: Precision of methods Step 1 (feature selection). True positive count among selected mediators, representing the number of correctly identified causal CpGs among the top-$m_{\mathbf{causal}}$ ranked by $|\alpha\beta|$ for HIMA and $\max\{P_\alpha, P_\beta\}^2$ for HDMAX2-surv. Simulation parameters: $m_{\mathbf{causal}} = 100$ causal probes among $p = 10000$ total CpGs, $\mu_\beta \in \{0.05, 0.5, 5\}$, $\sigma_\beta = 0.1$. Exposure $\rightarrow$ mediator effects: $\mu_\alpha = 0$ (no exposure $\rightarrow$ mediator effect), $\sigma_\alpha = 0$. $\pi_{\mathbf{overlap}} = 0.8$ Two sample sizes are shown ($n = 150$, purple; $n = 300$, orange) across 5 replications.**

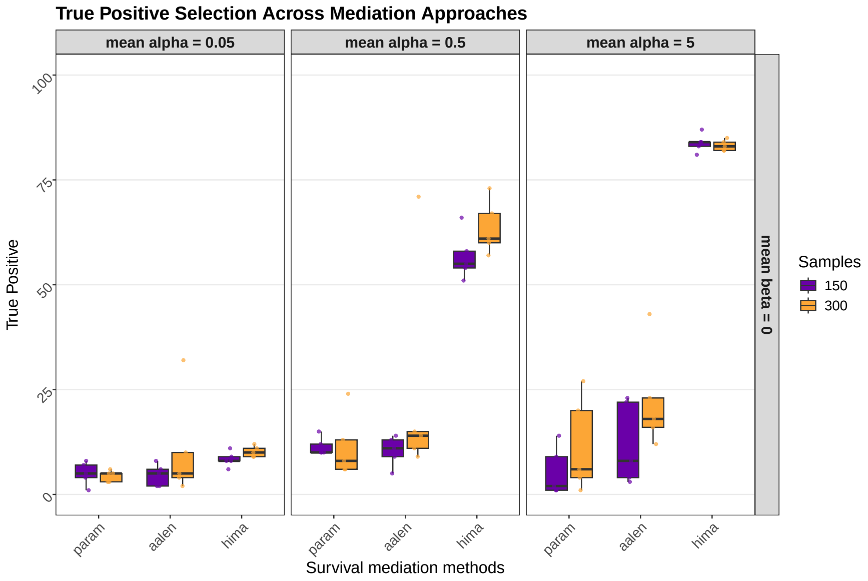


**Supplementary Figure 6: Precision of methods Step 1 (feature selection). True positive count among selected mediators, representing the number of correctly identified causal CpGs among the top-$m_{\mathbf{causal}}$ ranked by $|\alpha\beta|$ for HIMA and $\max\{P_\alpha, P_\beta\}^2$ for HDMAX2-surv. Simulation parameters: $m_{\mathbf{causal}} = 100$ causal probes among $p = 10000$ total CpGs, $\mu_\beta = 0$ (no mediator $\rightarrow$ outcome effect), $\sigma_\beta = 0$. Exposure $\rightarrow$ mediator effects: $\mu_\alpha \in \{0.05, 0.5, 5\}$, $\sigma_\alpha = 0.1$. $\pi_{\mathbf{overlap}} = 0.8$ Two sample sizes are shown ($n = 150$, purple; $n = 300$, orange) across 5 replications.**

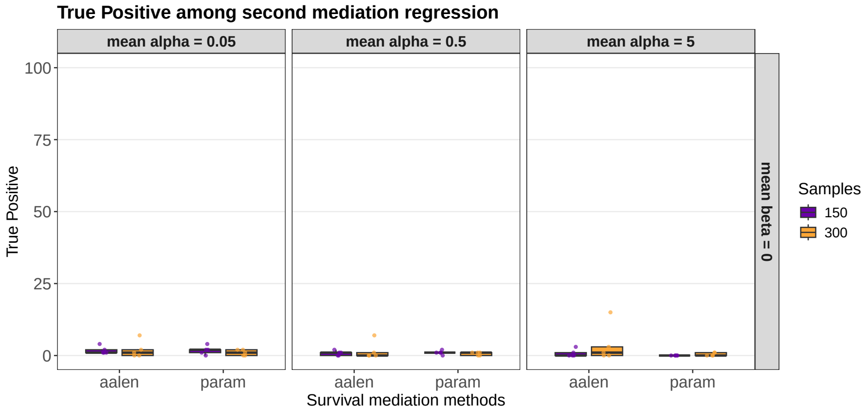


**Supplementary Figure 7: Precision of mediator → outcome regression ($\beta$ pathway). True positive count among selected mediators, representing the number of correctly identified causal CpGs among the top-$m_{\textbf{causal}}$ ranked by $P_\beta$ values. Simulation parameters: $m_{\textbf{causal}} = 100$ causal probes among $p = 10000$ total CpGs, $\mu_\beta = 0$ (no mediator → outcome effect), $\sigma_\beta = 0$. Exposure → mediator effects: $\mu_\alpha \in \{0.05, 0.5, 5\}$, $\sigma_\alpha = 0.1$. $\pi_{\textbf{overlap}} = 0.8$ Two sample sizes are shown ($n = 150$, purple; $n = 300$, orange) across 5 replications. HIMA is not included because this information cannot be extracted from its method implementation. These results confirm that the false positives observed for HDMAX2-surv in Supplementary Figure 6 are attributable to the $\max\{P_\alpha, P_\beta\}^2$ testing procedure used in the Step 1 selection stage.**

### 3 Mediation of smoking status on PDAC survival

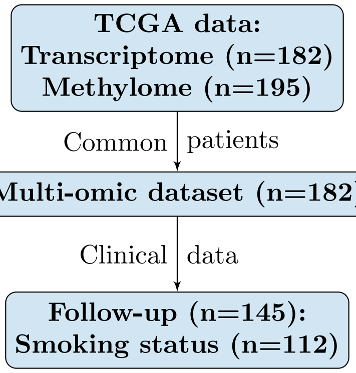


**Supplementary Figure 8: Filtering of the TCGA dataset to obtain a complete multi-omic dataset with follow-up information. The $n$ indicates the number of samples, with some patients having several samples.**

#### 3.1 Cohort selection

The Cancer Genome Atlas (TCGA) is the result of a collaboration between the National Cancer Institute (NCI) and the National Human Genome Research Institute (NHGRI) to generate molecular maps in major types and subtypes of cancer. It includes omic and clinical data on enrolled cases provided by Biospecimen Core Repositories (BCRs) in schema-valid XML format. We downloaded the TCGA-PAAD study which is the PDAC atlas.

| **Variables** | n (%) |
|---|---|
| Cohort | 112 (100%) |
| Age | 65 (Q1=57, Q3=74) |
| Sex | |
| Female | 55 (49%) |
| Male | 57(51%) |
| Tumor grade | |
| I | 5 (4.4%) |
| II | 50 (44.6%) |
| III | 56 (50%) |
| IV | 1 (0.8%) |
| Smoking | |
| Smoker (a and b) | 37 (33%) |
| (a) Current Smoker | 16 (14%) |
| (b) Reformed Smoker for $\leq$ 15 yrs) | 21 (19%) |
| Non smoker (c and d) | 75(67%) |
| (c) Reformed Smoker for $>$ 15 yrs | 24 (21%) |
| (d) Lifelong Non-Smoker | 51 (46%) |
| Deaths | 59 (53%) |

**Supplementary Table 3: Descriptive characteristics of the TCGA-PAAD cohort (n = 112). Age is reported as median with first and third quartiles (Q1–Q3). Sex, tumor grade, smoking status, and number of deaths are presented as counts and percentages.**

###### 3.1.1 Co-variables

We also included co-variables in the models, such as age, gender, and grade of the tumor. In PDAC histologic grade is based on gland-forming differentiation and is reported as G1 well differentiated, G2 moderately differentiated, G3 poorly differentiated, and G4 undifferentiated/anaplastic.

###### 3.1.2 Molecular data

We used transcriptomic and methylation data of PDAC patients from the TCGA (n=184 patients with methylation data, n=178 for RNAseq) filtered as in (8). The processing of the data was performed by the TCGA. For the methylation dataset, we also filtered out probes with NA values and probes located on sex chromosomes, retaining 371,033 features.

###### 3.1.3 Exposure assessment

Among the tobacco-related variables available in the TCGA database, we used `exposures.tobacco_smoking_status`, which includes six categories: "Lifelong Non-Smoker", "Current Reformed Smoker for $>$ 15 yrs", "Current Reformed Smoker for $\leq$ 15 yrs", "Current Smoker", "Current Reformed Smoker, Duration Not Specified", and "Unknown".

Patients with an "Unknown" smoking status ($n = 36$) were excluded from the analysis due to missing information. Patients categorized as "Current Reformed Smoker, Duration Not Specified" ($n = 8$) were excluded, as their smoking status could not be reliably assigned to either group.

We then created a binary smoking variable: "Lifelong Non-Smoker" and "Current Reformed Smoker for $>$ 15 yrs" were classified as **non-smokers** (0), while "Current Reformed Smoker for $\leq$ 15 yrs" and "Current Smoker" were classified as **smokers** (1).

###### 3.1.4 Follow-up

The follow-up variables are described as follows in the GDC Data Dictionary: "A visit by a patient or study participant to a medical professional. A clinical encounter that encompasses planned and unplanned trial interventions, procedures, and assessments that may be performed on a subject. A visit has a start and an end, each described with a rule. The process by which information about the health status of an individual is obtained before and after a study has officially closed; an activity that continues something that has already begun or that repeats something that has already been done." We used the following follow-up variables: (i) follow-up time (in days) and (ii) status (Censored = 0, event (death) = 1). We kept only common samples for which we have smoking, follow-up information, and non-zero survival time (n=112) (Table 3).

models, such as age, gender, and grade of the tumor. In PDAC histologic grade is based on gland-forming differentiation and is reported as G1 well differentiated, G2 moderately differentiated, G3 poorly differentiated, and G4 undifferentiated/anaplastic.

##### 3.2 Execution of HDMAX2-surv on the TCGA-PAAD cohort

###### 3.2.1 `HDMAX2-surv_step1`.

We ran `HDMAX2-surv.aalen` with cohort, age at diagnosis, gender, and grade as observed confounding factors, and we estimate $K$ unobserved latent factor. The optimal $K$ had been estimated using Cattell's rule applied to the eigenvalues of principal component analysis (PCA) (9). In the TCGA cohort, $K$ was estimated to be 8 based on PCA of the residuals obtained after regressing methylation values in the adjustments factors (Supplementary Figure 9).

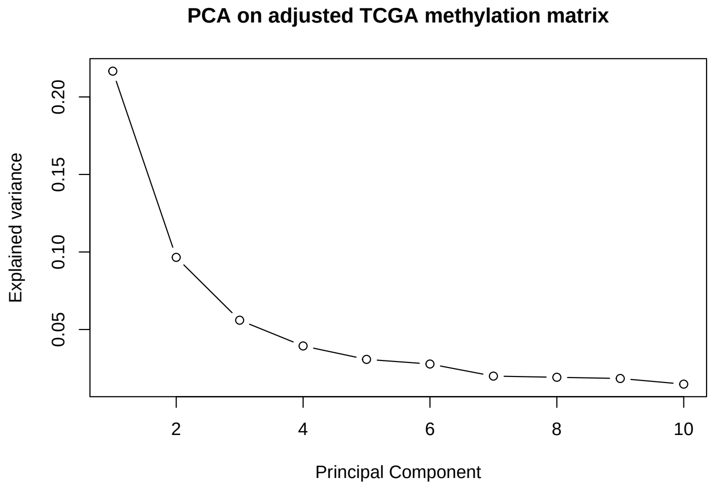


**Supplementary Figure 9: TCGA dataset PCA on methylation matrix adjusted on covariates screeplot).**

The Latent Factor were estimated using a Latent Factor Mixed Model (LFMM) (10) :

$$M = T\alpha + \text{Imm}\,\beta^{\top} + \text{Cov}\,\gamma^{\top} + W + E \tag{1}$$

with $M$: the methylation matrix, $T$: a vector of tobacco exposure, Imm: the immune cell-type composition matrix (estimated by deconvolution), Cov: the matrix of covariates (gender, age, tumor grade), $W = UV^{\top}$:

the latent matrix, $E$: the residual noise. The $K$ latent factors represent unobserved confounders, modeled through the matrix $U$, which is obtained via a singular value decomposition (SVD) of the matrix $W$, where $V$ is the matrix of loadings. During the latent-factor estimation step, we incorporated immune-infiltration profiles, derived from the deconvolution pipeline (see section 4.1), as fixed-effect covariates in the LFMM, thereby estimating latent factors conditional on these profiles. In subsequent analyses, we did not carry these immune-infiltration profiles forward: the high-dimensional mediation models were fitted without including them as covariates.

We then ran the function `run_AS_surv_param()` to evaluate the association between tobacco exposure, DNA methylation patterns, and survival outcomes, using the optimal number of latent factors $K$, previously determined, and the variables age at diagnosis, gender, and grade as observed adjustment factors.

###### 3.2.2 AMR detection

We used a p-value aggregation approach to identify Aggregated Methylated Regions (AMR) of interest. We used `comb-p` (11; 12), a method that combines adjacent CpG p-values in sliding windows, as described in Jumentier et *al* (5).

The COMBP method (12) identifies genomic regions enriched for statistically significant signals by locally combining p-values from individual tests at probes or genomic positions. This approach is particularly well suited to data with spatial dependence.

Its theoretical basis is the Stouffer–Lipták p-value combination test, which integrates information from multiple (correlated) tests:

$$Z = \frac{\sum_{i=1}^{k} w_i \Phi^{-1}(1 - p_i)}{\sqrt{\sum_{i=1}^{k} w_i^2}}$$

Here, $Z$ is the combined score; $p_i$ is the p-value from test $i$; $\Phi^{-1}$ is the quantile function of the standard normal distribution; $w_i$ is the weight assigned to test $i$; and $k$ is the total number of tests considered.

Under the null hypothesis, the combined $Z$ follows a standard normal distribution and can be converted to an overall p-value. In genomic data, p-values are often autocorrelated because probes are physically close. The COMBP procedure accounts for this dependence by estimating the empirical autocorrelation of the $P$-values via the autocorrelation function (ACF). Regions containing several consecutive positions with significant combined p-values are called enriched regions. Each region is then assigned an overall p-value obtained by combining the individual p-values it contains.

In the original HDMAX2 implementation of Jumentier et al. (2022) (5), COMBP was adapted for methylation probe data and for the detection of AMRs—aggregates of signals—rather than DMRs (Differentially Methylated Regions), which represent mean methylation differences between groups. We chose to retain this adaptation as is. We defined significant regions as those containing at least two markers at a maximum distance of 1,000 bp and significant at the 5% threshold. The requirement of at least two CpGs within 1 kb ensures spatial coherence and limits isolated probe artefacts, consistent with established DMR detection practices. CpGs located within 1,000 bp are typically highly correlated due to shared regulatory context, supporting regional aggregation. Averaging across CpGs provides a robust summary of the regional methylation signal, reduces probe-level noise, and stabilizes high-dimensional mediation estimation.

###### 3.2.3 `HDMAX2-surv_step2`.

In the second stage, we evaluated two scenarios. In Scenario A, we treated individual CpG sites as mediators and declared significance under false discovery rate control at 5% and 10% (Supplementary Figure 10).

In Scenario B, we first constructed AMRs and carry forward the 50 most significant AMRs from that step. We then applied the `estimate_effect_surv_param()` function to evaluate the indirect effects of AMRs in the pathway between tobacco exposure and survival, using the approach implemented in the `mediation` R package. We selected 36 significant AMRs with $P-value \leq 0.05$, which corresponds to adjusted $P-value \leq 0.065$ after Benjamini-Hochberg correction for multiple testing using the R function `p.adjust()` with `method = "BH"` Supplementary Table 4).

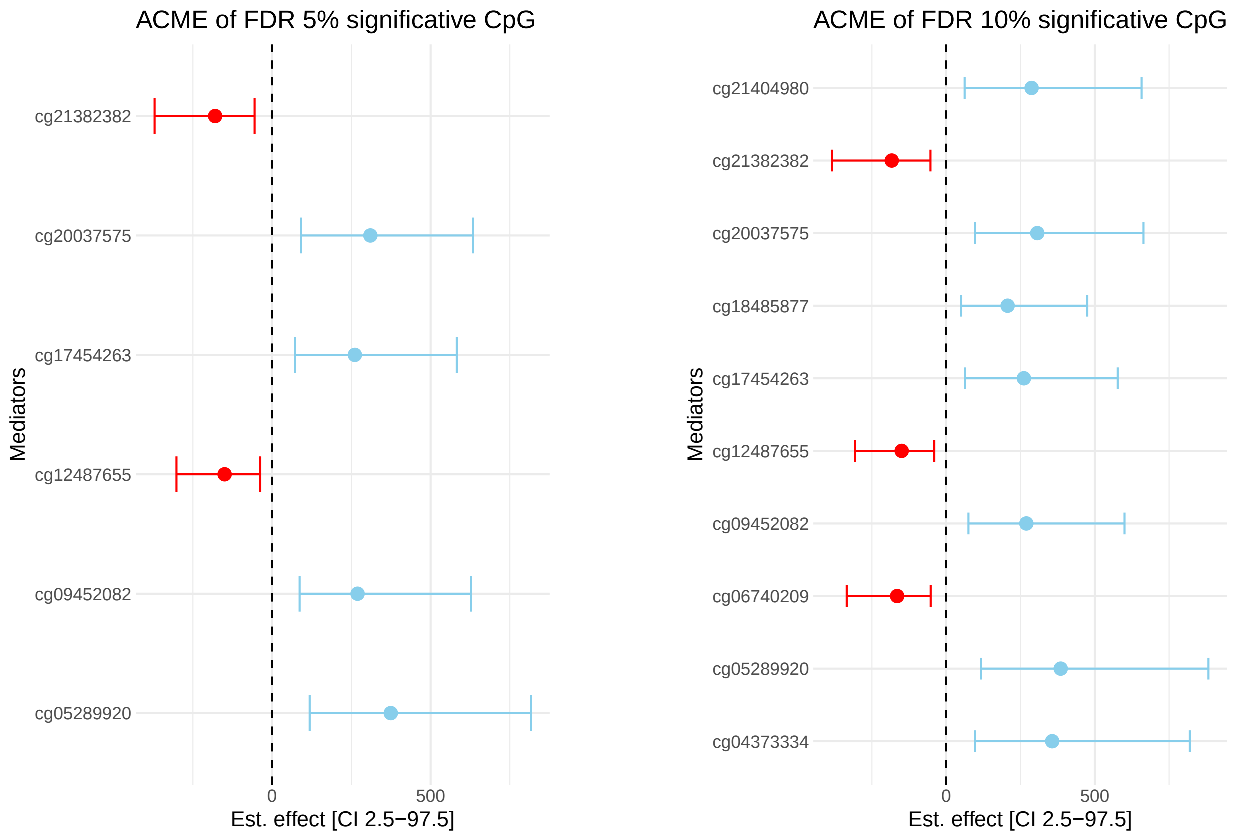


**Supplementary Figure 10: Average Causal Mediation Effects (ACMEs) of DNAm CpGs with 5% and 10% FDR significance from TCGA-PAAD tobacco to survival mediation analysis. Several CpGs were also captured by the AMR-based approach, including cg21382382 (AMR38), cg20037575, cg09452082, and cg05289920 (AMR1), as well as cg04373334 (AMR23).**

##### 3.2.4 Analysis of four candidate genes

Among the significant AMRs, we identified four genes previously associated with PDAC occurrence or outcome, defined as having more than two prior citations in PubMed (Supplementary Figure 11).

*MSX2*-AMR2, a member of the homeobox gene family, a target of the RAS signaling pathway, expressed in pancreatic cancer and related to chemoresistance (13; 14). *KCNQ1*-AMR5, a potassium channel gene subject to tissue-specific imprinting and found to be differentially methylated in pancreatic cancer (15); *PIWIL1*-AMR17, a member of the Argonaute protein family, implicated in pancreatic fibrosis and the regulation of pancreatic cancer (16; 17); and *EPHA3*-AMR37, part of the ephrin receptor family, which contributes to glucose homeostasis in insulin-producing pancreatic cells (18)

Among the significant AMRs, we identified four genes previously associated with PDAC occurrence or outcome, defined as having more than two prior citations in PubMed (Supplementary Figure 11).

AMR2 exhibited an ACME of 283.7 [95% CI: 30.6, 830.0] and AMR37 an ACME of 187.1 [95% CI: 1.9, 502.2]. In both cases, tobacco exposure globally increased methylation at the locus in smokers compared to non-smokers. This hypermethylation shifts the survival curve forward, thereby improving survival through a protective mediation pathway with synergistic positive effects.

In contrast, AMR5 (123.6 [95% CI: 19.3, 299.3]) and AMR17 (87.7 [95% CI: 1.2,235.3]) showed the opposite trend: tobacco exposure decreased methylation, and methylation at these loci was negatively associated with survival. Consequently, hypomethylation conferred an overall protective effect, mediated through synergistic negative effects.

| AMR | Chromosome | Size (pb) | CpGs | Genes | ACME_CI | P-value | Pubmed Hits |
|---|---|---|---|---|---|---|---|
| AMR1 | chr15 | 2906 | cg00032912, cg00033666, cg00436496, cg02326806, cg05289920, cg07116712, cg07959469, cg09452082, cg10151901, cg10186131, cg17140992, cg17454263, cg18761422, cg20037575, cg22460896, cg22703162, cg22840361, cg24491784 | RP11-262 | 410.6 [117.2,976.1] | 0.000 | 0 |
| AMR2 | chr5 | 1321 | cg01960016, cg02365742, cg06946371, cg12387713, cg15123984, cg17156565, cg20563910, cg25998402, cg26615160, cg26796283, cg27096144 | MSX2 | 283.7 [30.6,830.0] | 0.012 | 9 |
| AMR4 | chr14 | 683 | cg02814525, cg09255505, cg14163691, cg14918743, cg19515081 | ZBTB42 | 264.6 [38.8,687.4] | 0.004 | 0 |
| AMR5 | chr11 | 670 | cg00283576, cg01178971, cg03371125, cg04104132, cg04902871, cg10678459, cg11025829, cg11363972, cg15932136, cg19698309, cg19779211, cg20533553, cg23267890, cg23623667, cg24079038 | KCNQ1 | 123.6 [19.3,299.3] | 0.014 | 4 |
| AMR7 | chr17 | 299 | cg10997810, cg15229994, cg16293484, cg19948701, cg20030711 | SEPT9 | 130.1 [11.6,342.4] | 0.028 | 1 |
| AMR8 | chr19 | 441 | cg02587316, cg04332534, cg05020604, cg18630667, cg22517995, cg25397945 | ZNF382 | 119.7 [4.2,334.8] | 0.040 | 2, 0 |
| AMR9 | chr4 | 609 | cg00169305, cg01725318, cg02651961, cg03168293, cg03575792, cg03710295, cg08129449, cg08821557, cg11539424, cg21246783, cg24695316, cg25323711 | CLGN | 231.7 [23.6,669.4] | 0.020 | 0 |
| AMR10 | chr7 | 680 | cg09016797, cg12712184, cg13104880, cg14635659, cg14938587, cg21428990 | | 174.2 [37.4,416.3] | 0.008 | |
| AMR11 | chr7 | 359 | cg00702008, cg06129054, cg06994813, cg13909585, cg15478390, cg17137182, cg17611951, cg24399959 | PCLO | 93.7 [1.1,285.2] | 0.046 | 1 |
| AMR12 | chr20 | 219 | cg06198069, cg11252765, cg22582721 | BLCAP, NNAT | 272.3 [50.7,640.7] | 0.006 | 0, 0 |
| AMR13 | chr10 | 879 | cg02676262, cg05477444, cg15847996, cg24571871 | INPP5A | 109.3 [8.3,288.6] | 0.024 | 0 |
| AMR15 | chr6 | 553 | cg01350077, cg01883425, cg05345286, cg06688989, cg27200446 | MDFI | 160.3 [25.2,403.8] | 0.008 | 2 |
| AMR17 | chr12 | 155 | cg06365016, cg12020444, cg16244664, cg27261050 | PIWIL1 | 87.7 [1.2,235.3] | 0.044 | 4 |
| AMR18 | chr7 | 288 | cg05767159, cg17400476 | RP1-170O | 185.2 [38.1,410.7] | 0.002 | 0 |
| AMR19 | chr1 | 342 | cg04486940, cg14210311, cg16681083, cg26233866 | ERI3 | 116.7 [15.2,297.4] | 0.016 | 0 |
| AMR20 | chr2 | 285 | cg03002352, cg04804377, cg10167837, cg15124757, cg21642176, cg25565383, cg26119367 | RAD51AP2 | 114.7 [0.2,368.8] | 0.050 | 0 |
| AMR21 | chr19 | 227 | cg03657031, cg06697294, cg11662224, cg21950287, cg23460210, cg27552287 | PRKCG | 80.7 [0.5,216.5] | 0.046 | 1 |
| AMR22 | chr14 | 155 | cg02054667, cg11051738 | | 152.5 [23.7,402.9] | 0.010 | |
| AMR23 | chr4 | 92 | cg04373334, cg08575049 | FAT4 | 143.9 [21.7,355.6] | 0.006 | 1 |
| AMR25 | chr17 | 619 | cg00117908, cg07004514, cg08901098, cg09980477, cg11102858, cg13173111, cg15253816, cg25456728 | ARSG | -114.3 [-260.1,-7.3] | 0.032 | 1 |
| AMR26 | chr16 | 286 | cg02612270, cg06430465, cg10244976, cg13706784 | LMF1 | 115.2 [4.2,290.4] | 0.032 | 0 |
| AMR27 | chr1 | 227 | cg01101873, cg06060874, cg27187555 | PRDM16 | 165.7 [24.5,396.5] | 0.010 | 1 |
| AMR29 | chr10 | 106 | cg02987635, cg14207539, cg23264016 | C10orf11 | 128.8 [12.2,354.3] | 0.018 | 0 |
| AMR30 | chr1 | 181 | cg07533148, cg20146541, cg20810478, cg23054189, cg26157385 | TRIM58 | 107.2 [3.3,297.9] | 0.034 | 1 |
| AMR33 | chr3 | 131 | cg08002883, cg10511904, cg16376000, cg17393267 | FGF12 | 103.6 [0.4,258.1] | 0.048 | 0 |
| AMR35 | chr19 | 253 | cg01174276, cg02671333, cg12029794, cg23854456 | CTC-246B | -114.0 [-260.3,-19.4] | 0.006 | 0, 0 |
| AMR36 | chr2 | 388 | cg02304863, cg02518161, cg06870284, cg08194677, cg12784598, cg13165472, cg16675700, cg18564808 | ANTXR1 | 101.0 [1.5,308.7] | 0.046 | 0 |
| AMR37 | chr3 | 353 | cg02517134, cg05676541, cg10315231, cg18055394, cg22619563 | EPHA3 | 187.1 [1.9,502.2] | 0.050 | 3 |
| AMR38 | chr4 | 332 | cg10024909, cg15234400, cg16919569, cg21382382 | HAND2-AS | -123.5 [-285.1,-6.4] | 0.040 | |
| AMR39 | chr1 | 429 | cg03199058, cg06802365, cg10174867, cg11804724, cg20492807, cg25379026 | DNAH14 | 105.5 [0.9,309.3] | 0.048 | 0 |
| AMR40 | chr5 | 151 | cg04223420, cg07274716, cg22827250 | PITX1 | 129.5 [15.2,347.1] | 0.006 | 1 |
| AMR41 | chr8 | 381 | cg03979241, cg04662594, cg14837598, cg16678564, cg26392737 | DMTN | 91.0 [1.4,264.6] | 0.048 | 0 |
| AMR46 | chr14 | 210 | cg02005147, cg02867079, cg03475665, cg06834875, cg06969206, cg09956302, cg12188268, cg26669257 | HHIPL1 | -79.7 [-220.5,-1.6] | 0.042 | 0 |
| AMR47 | chr8 | 98 | cg01906015, cg06410746, cg08855288, cg15858239 | NKAIN3 | 165.9 [2.8,506.9] | 0.038 | 0 |
| AMR48 | chr12 | 472 | cg01802493, cg05863637, cg06153623, cg16236851, cg17226286 | RP11-474 | -88.6 [-207.5,-9.0] | 0.036 | 0, 0 |
| AMR49 | chr6 | 111 | cg17284168, cg18566177, cg21366688 | SGK1 | 130.1 [4.1,356.1] | 0.024 | 2 |

**Supplementary Table 4: Significant mediator-associated methylation regions (AMRs). Each AMR is defined by its chromosomal location, size (in base pairs), included CpG sites, annotated genes, the estimated Average Causal Mediation Effect (ACME) with 95% confidence interval, and the associated $p$-value. The "Pubmed Hits" column shows the number of PubMed articles retrieved with the query: *Humans[Mesh] AND (pancreatic cancer[MeSH]) AND GENE*. Empty cells indicate missing gene annotation or no publication found.**

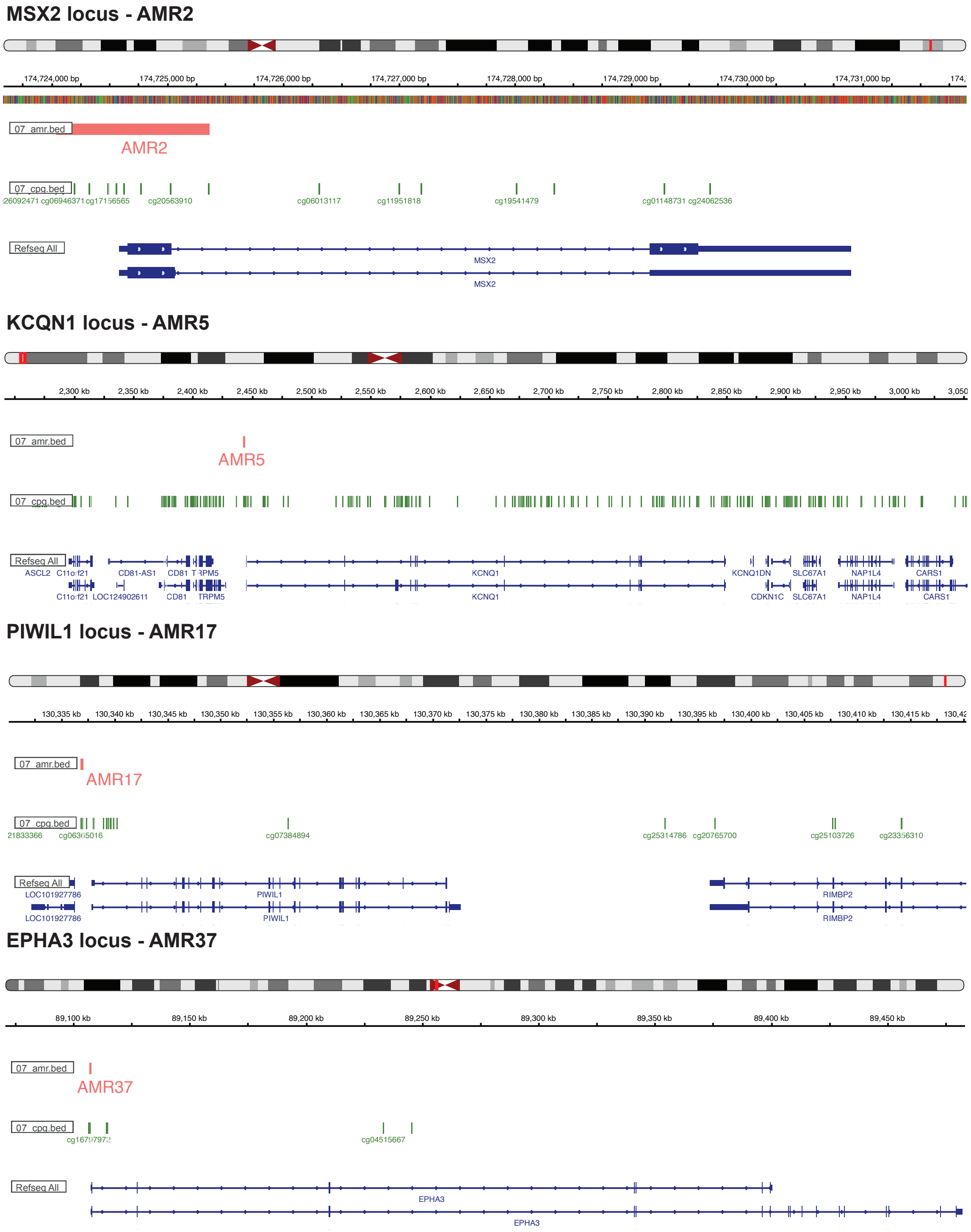


**Supplementary Figure 11: IGV representation of the locus of know genes proximal to significant AMRs.**

### 4 Causal discovery of tobacco-survival mechanisms

#### 4.1 Quantification of immune infiltration on the TCGA-PAAD cohort

##### 4.1.1 Reference profiles

To perform supervised deconvolution, we used a dataset of bulk transcriptomic profiles from nine pure cell types found in PDAC: endothelial cells, neutrophils, macrophages, B cells, $CD4^{+}$ and $CD8^{+}$ cells, fibroblasts, and classical and basal-like tumour cells (GSE281204). We also downloaded two published single-cell datasets with labels to use as single-cell references during deconvolution. The first dataset (19) contained 57,530 cells with the following cell types labelled by the authors: fibroblasts, stellate cells (renamed fibroblasts), macrophages, endothelial cells, T and B cells, ductal cells type 2 (renamed tumor cells) and 1, endocrine cells, and acinar cells. The second dataset (20) contained 66,075 cells with the following cell types: macrophages, NK cells, regulatory T cells, tumor cells, B cells, hepatocytes, pDCs and DCs, endothelial cells, plasma cells, and mesenchymal cells. We retained in both datasets, respectively 44,549 and 22,662 cells that match the cell types present in the bulk reference (endothelial cells, B and T cells, macrophages, fibroblasts, tumor cells for the Peng dataset (19), endothelial cells, B cells, macrophages, NK cells, regulatory T cells, pDCs and DCs and tumor cells for Raghavan (20)).

For each single-cell dataset, we sampled 5,000 cells randomly (to reduce computing time) while retaining the same number of cells from each patient (n=35 for Peng, n=2 for Raghavan), and used puriST, a classifier that separates tumor cells into basal or classical-like (21).

##### 4.1.2 Deconvolution

We applied supervised deconvolution algorithms to retrieve cell type proportions (see Supplementary Figure 12). We deconvoluted the transcriptomic profiles of the samples used in the mediation analysis, using bulk or single-cell data as references (GSE281204 for the bulk reference, (19) and (20)) for the single-cell ones). We applied robust linear regression (RLR) with bulk references, and SCDC (22), InstaPrism (23), and MuSiC (24) with single-cell references. For RLR, we used the function `epidish` from the package EpiDISH with the "method" parameter set to "RPC." Of note, the references were normalized with DESeq2 beforehand (25). For InstaPrism (23), we used the function `InstaPrism` from the eponymous package. We got two proportions matrices, one per single-cell reference. For SCDC (22), we used the function `SCDC_prop` to estimate the proportion matrix from a single reference. For MuSiC (24), we used the function `music_prop from` the package MuSiC and obtained a proportion matrix per single-cell reference. We finally computed a consensus prediction, based on the 7 predictions from the four methods described above. The consensus prediction was done based on the following procedure. First, all subsets of dendritic cells (DCs) and T cells (resp. plasmacytoid and $XCR1^{+}$ DCs for DCs and $CD4^{+}$, $CD8^{+}$ and regulatory T cells for T cells) were aggregated into a single type, respectively, called DCs and T cells. For the predicted proportion matrix, we sumed the proportions of all DCs (resp. T cells) subsets into a single type of DCs (resp. T cells) ($A_{\text{avg}}$). For the bulk reference matrix, we computed the average profile for DCs and T cells. For single-cell references, we first performed a pseudo-bulk before averaging the reference profiles of the DCs and T cell subsets ($T_{\text{avg}}$). Then for each prediction, we computed the Root Mean Square Error (RMSE) between the reconstruction $T_{\text{avg}} \times A_{\text{avg}}$ and the expression matrix. Finally, we computed the consensus proportion for each cell type. If a cell type is estimated by a single method out of the 7 described above, the consensus proportion is the prediction of this method. Otherwise, we performed a weighted average of all methods predicting this cell type, the weights being the normalized inverse of the RMSEs.

This consensus approach allowed us to quantify the composition of tumor microenvironment across 9 distinct cellular components, including 5 immune cell types: B cells, T cells, dendritic cells (DCs), natural killer (NK) cells, and macrophages.

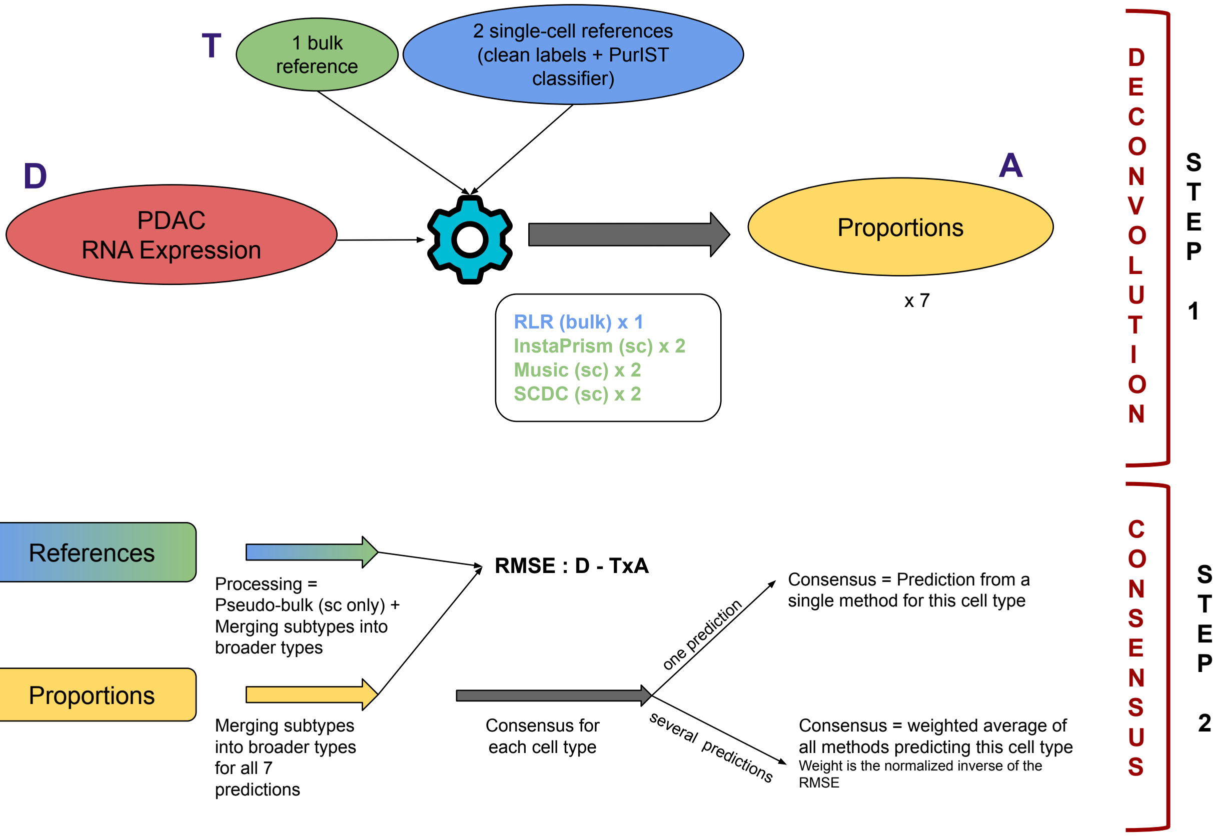


**Supplementary Figure 12: Flowchart of the deconvolution pipeline using transcriptomic data and consensus references/methods**

#### 4.2 Causal discovery framework for immune-mediated pathways

We developed a causal discovery framework to identify the relationships between tobacco exposure, DNA methylation, and immune infiltration in the survival of PDAC patients. We investigated the causal relationships between tobacco exposure and patient survival through immune-mediated pathways using a directed acyclic graph (DAG) framework. The analysis included six key variables: tobacco exposure status ($T$, binary: smoker vs. non-smoker), DNA methylation levels of significant AMRs ($A$, continuous), total immune infiltration proportion ($I_{all}$), B cell infiltration proportion ($I_B$, continuous), T cell infiltration proportion ($I_T$, continuous), and survival outcome ($S$, continuous, measured as accelerated failure time). We first started from a fully connected graph (Supplementary Figure 13) and then assessed sequentially the existence and directionality of each node.

##### 4.2.1 Exposure to tobacco $T$

Starting from a fully connected graph, we tested unconditional independence between exposure to tobacco ($T$) and survival ($S$). Using a cox proportional hazards model ($P$-value = 0.80760), we identified no significant association between tobacco ($T$) and survival ($S$). The model was adjusted for observed confounding factors: age at diagnosis, sex, and tumor grade.

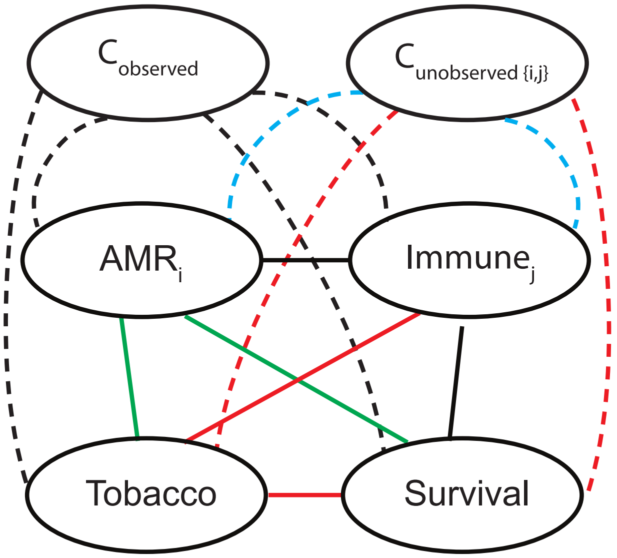


**Supplementary Figure 13: Complete directed acyclic graph (DAG) framework.**
**Observed confounders (age, sex, and tumor grade) are denoted $C_O$, latent factors (LFs) representing potential unobserved confounders are denoted $C_U$, tobacco exposure status is denoted $T$, AMRs are denoted $A_i$ , and immune fractions are denoted $I_j$.**
**Red edges indicate relationships removed by unconditional independence testing: section 4.2.1 − $S \perp T|C_O$: section 4.2.2 − $C_U \perp T$ and $C_U \perp S$; section 4.2.3 − $T \perp I_j$ where $j \in \{\textbf{Tot.Imm.}, \textbf{B cells}, \textbf{T cells}\}$.**
**Green edges show relationships retained after testing HDMAX2-surv testing: $T \not\perp A_i$ and $A_i \not\perp S$ for $i \in \{1, 2, \ldots, 36\}$.**
**Blue edges represent latent factor confounding relationships that are specific to individual AMR-immune pair combinations (see section 4.2.2).**

###### 4.2.2 Unobserved confounders $C_U$ : selection of the latent factors (LF)

We first tested that HDMAX2-surv LFs were uncorrelated with tobacco exposure, methylation, immune infiltration and survival. LFs ($C_U$) were originally estimated from the model $M\ T|C_O, I, C_U$. Because our objective was to construct a specific causal graph for each $A$-$I$ pair, we re-evaluated the dependence between LFs and each variable in the model separately for every pair. We used conservative Bonferroni correction for latent factor identification to minimize model complexity and preserve statistical power.

First, we tested the dependence between tobacco exposure and each of the eight LFs using Pearson correlation coefficients. All latent factors showed non-significant correlations with tobacco exposure (Pearson correlation $P$-values: 0.8216, 0.8814, 0.4849, 0.2818, 0.8056, 0.8428, 0.0891, and 0.3950 for LFs A to H, respectively).

Second, we tested the dependence of LFs with patient survival (Cox model $P$-values: 0.9403, 0.6609, 0.7011, 0.0619, 0.4923, 0.3478, 0.4997, and 0.6096 for LFs A to H, respectively).

Then, to ensure the validity of our causal framework and avoid spurious confounding relationships, we systematically evaluated the associations between HDMAX2-surv-derived latent factors (LFs, 8 components of the matrix $U$) and both immune infiltration variables and AMRs. This analysis aimed to identify latent factors that could represent unmeasured confounders in the causal pathways under investigation.

We computed Pearson correlation coefficients between all latent factors ($n = 8$) and both immune infiltration variables ($n = 6$: total immune, B cells, T cells, DCs, NK cells, and macrophages) and AMRs ($n = 50$) (Supplementary Figure 14). Statistical significance was assessed using Pearson correlation tests ($P - value < 0.01$), with a Bonferroni correction for multiple testing. To identify latent factors that could act as confounders in AMR-immune relationships, we systematically examined all possible AMR-immune pairs ($n = 50 \times 6$ combinations). For each pair, we identified latent factors that simultaneously exhibited significant correlation with both the AMR and the immune variable. Such latent factors were flagged as potential confounders and accounted as covariates in the later models.

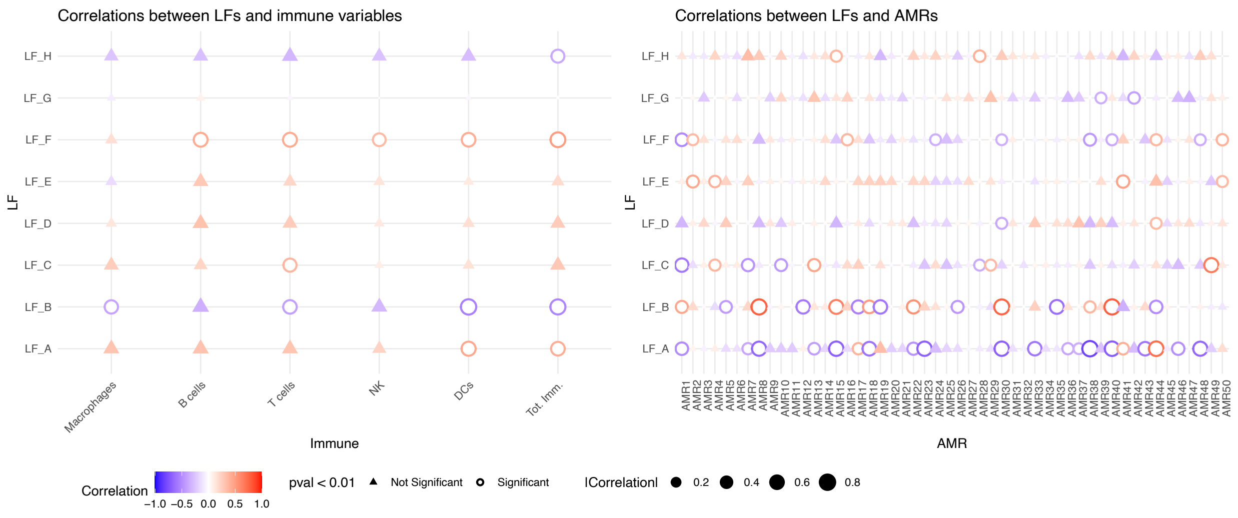


**Supplementary Figure 14: Correlation of Latent Factors with AMR and Immune variables. Correlation patterns are visualized using dot plots where point size represented correlation magnitude, color indicated correlation direction (blue for negative, red for positive), and shape distinguished between statistically significant and non-significant associations after Bonferroni correction.**

Of note, immune infiltration profiles ($I$) were included as conditioning variables only during the estimation of latent factors to capture high-dimensional residual variation from the methylation matrix. This step was purely statistical, aiming to extract latent sources of unwanted variation orthogonal to the exposure, while accounting for major structured biological signals. In contrast, in the mediation framework, immune infiltration is explicitly treated as a potential mediator in the serial pathway $T \to A \to I \to S$. For each AMR–immune pair analyzed, we systematically evaluate whether the immune variable acts as a confounder or mediator in the $A \to S$ pathway, ensuring unbiased estimation of causal effects.

##### 4.2.3 Immune variable $I$: survival anlaysis to select immune variables of interest

We implemented a two-tiered approach for causal discovery. In the following steps, given the exploratory nature of the causal discovery framework, we used a liberal significance threshold ($P < 0.15$) for conditional independence tests to maximize the sensitivity in identifying potential causal relationships. The final causal inferences were interpreted within the context of biological plausibility and effect size magnitudes.

**Dependance between Immune variables and survival.** Cox proportional hazards modeling revealed significant associations the 5 immune cell types (B cells, T cells, dendritic cells (DCs), natural killer (NK) cells, and macrophages) and patient survival outcomes (see Kaplan-Meier curves in Supplementary Figure 15). We identified 3 immune component displaying an associaion with survival: (i) total immune infiltration (sum of all five immune components; HR = 0.85 per 1% increase in immune fraction [95% CI: 0.69, 1.056], $P$-value = 0.1442), (ii) B cells (HR = 0.48 per 1% increase in B cell fraction [95% CI: 0.19, 1.22], $P$-value = 0.1229), and (iii) T cells (HR = 0.58 per 1% increase in T cell fraction [95% CI: 0.28, 1.19], $P$-value = 0.1402) .

All models were adjusted for observed confounding factors: age at diagnosis, sex, and tumor grade. These findings suggest that the composition of immune microenvironment significantly influences PDAC patient prognosis, which warrants further investigation of potential mediation mechanisms.

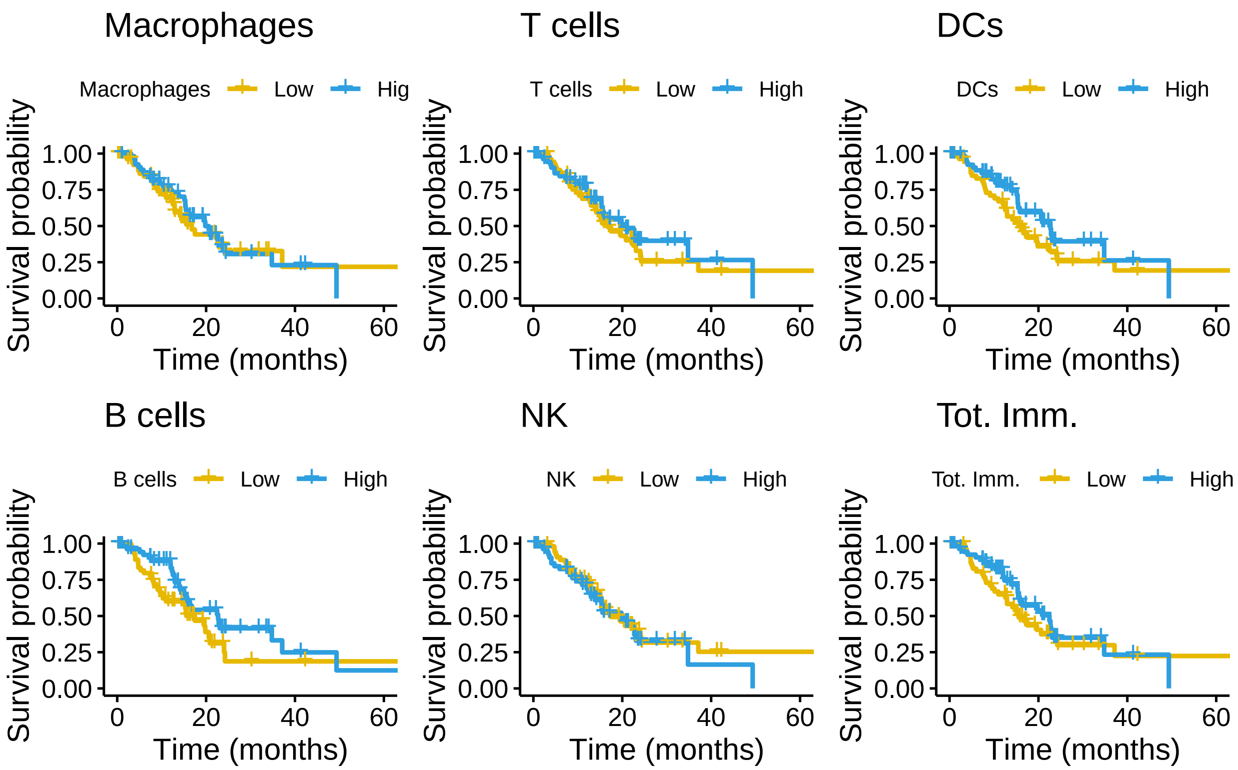


**Supplementary Figure 15: Kaplan–Meier survival compare overall survival between patients with low and high levels of the immune cell type. Immune infiltration was dichotomized based on median values for each cell-type. High expression is shown in blue, low expression in orange.**

**Independance between Immune variables and Tobacco.** Using Linear regressions, we tested for independance between Immune variables and Tobacco. We focused on immune variables linked with survival (total immune francion, B cells and T cells). All models were adjusted for observed confounding factors: age at diagnosis, sex, and tumor grade. We observed no association between theses three tobacco and Immune variables : $P_{\text{B cells}} = 0.96$, $P_{\text{T cells}} = 0.56$, and $P_{\text{Tot.Imm.}} = 0.51$)

###### 4.2.4 Systematic analysis of each AMR-immune $A$-$I$ pairs

To distinguish spurious associations from genuine causal relationships and determine the directionality of the causal graph, we combined biological prior knowledge (established mechanistic understanding) with conditional independence testing (statistical tests to identify dependencies and independencies among variables under different conditioning sets).

Based on biological prior knowledge and preliminary association analyses, we defined two candidate causal graphs (see Supplementary figure 16):

- **Graph 1**: $T \to A \to I \to S$ with additional direct effect $A \to S$
- **Graph 2**: $T \to A \to I \to S$ without direct effect $A \to S$

To determine which directed acyclic graph (DAG) was supported for each AMR-immune pair (including significant immune variables: $I_{all}$, $I_B$, and $I_T$), we employed a systematic edge-removal approach, starting from a fully connected graph (Supplementary Figure 13). First, we removed edges based on unconditional independence tests (all adjusted for age at diagnosis, sex, and tumor grade): (i) smoking and survival : $T \perp S$ (survival cox model, $P - value = 0.80760$); (ii) smoking and total immune infiltration: $T \perp I_{\text{Tot.Imm.}}$ (linear regression, $P - value = 0.513$); (iii) Smoking and B cells infiltration: $T \perp I_{\text{Bcells}}$ (linear regression,

$P-value = 0.954$); and (iv) smoking and Tcells infiltration: $T \perp I_{\mathrm{DCs}}$ (linear regression, $P-value = 0.564$). Finally, we filtered out AMRs for which significant associations were not confirmed with either smoking (linear model at $P > 0.15$) or survival (Cox model at $P > 0.15$). HDMAX2-derived Latent factors, identified as potential confounders for these specific pairs of AMR-immune, were included as additional nodes in the causal graph ($C_U$, in addition to the observed confounders $C_O$) (see **??** for more details). This systematic approach ensured that our causal inferences were not confounded by unmeasured factors captured by the latent factor model.

**Step 1: Unconditional independence testing.** Starting from a fully connected graph, we tested unconditional independence between all pairs of variables using appropriate statistical models. Although AMRs were previously identified as statistically significant mediators in the $T \rightarrow A \rightarrow S$ mediation analysis, the adjustment set differs for each AMR in the present framework, thereby justifying a dedicated unconditional independence testing step.

- Tobacco-AMR associations: Linear regression models. We filtered out AMRs that did not show significant associations with smoking ($P - value > 0.15$):

$$A = \alpha_0 + \alpha_T T + \boldsymbol{\alpha}_C^\top \mathbf{C} + \varepsilon \quad (2)$$

  where: $A$ denotes the methylation level (AMR), $T$ represents tobacco exposure (e.g., smoking status), and $\mathbf{C} = C_0 + \mathrm{LF}_s$ includes the observed covariates $C_0$ (age, sex, and tumor grade) together with the subset of $A$–$T$-specific significant latent factors ($\mathrm{LF}_s$) selected as described in Section 4.2.2. The error term $\varepsilon$ is assumed to follow a Gaussian distribution with constant variance.

- AMR-Survival associations: Cox proportional hazards models. We filtered out AMRs that did not show significant associations with survival ($P - value > 0.15$), as these AMRs would not contribute meaningfully to the mediation pathways under investigation.

$$\lambda_i(t \mid A, \mathbf{C}) = \lambda_0(t) \exp\Big(\beta_A A + \boldsymbol{\beta}_C^\top \mathbf{C_O}\Big) \quad (3)$$

  where: $\lambda(t \mid A, \mathbf{C_O})$ is the hazard function for each patient at time $t$, $\lambda_0(t)$ is the unspecified baseline hazard function, $A$ denotes the methylation level (AMR), and $\mathbf{C} = C_0 + \mathrm{LF}_s$ includes the observed covariates $C_0$ (age, sex, and tumor grade) together with the subset of $A$–$S$-specific significant latent factors ($\mathrm{LF}_s$) selected as described in Section 4.2.2.

**Step 2: Conditional independence testing.** To discriminate between Graph 1 and Graph 2, we tested whether AMRs retained a direct causal effect on survival after conditioning on immune infiltration. For each AMR-immune pair, we fitted the following cox model:

$$\lambda_i(t \mid A, I, \mathbf{C}) = \lambda_0(t) \exp\Big(\beta_A A + \beta_I I + \boldsymbol{\beta}_C^\top \mathbf{C}\Big) \quad (4)$$

where: $\lambda(t \mid M, \mathbf{C})$ is the hazard function for each patient at time $t$, $\lambda_0(t)$ is the unspecified baseline hazard function, $A$ denotes the methylation level (AMR), $I$ denotes the immune fraction, $\mathbf{C}$ represents the covariates (age, sex and grade) together with the subset of $A$–$I$-specific significant latent factors ($\mathrm{LF}_s$) selected as described in Section 4.2.2.

We retained only AMR-immune pairs where the immune variable showed significant association with survival ($P < 0.15$). The causal graph was determined based on the significance of the AMR coefficient ($\beta_A$):

- If $\beta_A$ remained significant ($P < 0.15$): Graph 1 (direct effect $A \rightarrow S$ preserved)
- If $\beta_A$ became non-significant ($P \geq 0.15$): Graph 2 (conditional independence $A \perp S \mid I$)

From there, we identified two possible candidate causal graphs: Graph 1, where AMRs have causal effects on both immune content and survival; and Graph 2, where AMRs have causal effects only on immune infiltration. Then we run conditional independence tests for graph discrimination (Supplementary Figure **??**D). To distinguish between Graph 1 and Graph 2 for each AMR-immune pair, we tested the presence of a direct edge from AMR to survival using the model 4. We retained only pairs in which the immune variable was significant ($P - value < 0.15$). If both $A$ and $I$ remained significant, we preserved the edge $A \to S$ (Supplementary Figure 16 Graph 1). When $A$ lost significance after controlling for the immune variable ($A \perp S \mid I$), we removed the edge $A \to S$ (Supplementary Figure 16 Graph 2).

**Step 3: Edge orientation.** Causal directions were established using biological prior knowledge for most edges:

- $T \to A$: Tobacco exposure precedes and causally affects DNA methylation
- $A \to S$ and $I \to S$: Molecular changes affect survival outcomes

In addition, as shown in Section 4.2.3, we found no evidence supporting a direct effect of tobacco exposure on immune infiltration $I \perp T | C_O$. To orient the edge between AMRs and immune infiltration ($A \leftrightarrow I$), we tested whether AMRs act as colliders by fitting the following logistic model:

$$\text{logit}\big(\Pr(T = 1 \mid A, I, \mathbf{C_O})\big) = \gamma_0 + \gamma_A A + \gamma_I I + \boldsymbol{\gamma}_C^\top \mathbf{C_O} \tag{5}$$

where: $T$ denotes tobacco exposure status(binary outcome), $A$ represents the methylation level (AMR), $I$ denotes the immune-related variable, $\mathbf{C_O}$ represents the covariates; $\gamma_0$ is the intercept, $\gamma_A$ and $\gamma_I$ are the log-odds ratios associated with AMR and immune infiltration, respectively, $\boldsymbol{\gamma}_C$ contains the regression coefficients for covariates. The edge was oriented as $A \to I$ when the immune variable remained conditionally independent of tobacco given the AMR ($T \perp I \mid A$, i.e., $\gamma_I$ non-significant at $P - value < 0.15$).

We retained pairs where the immune variable remained conditionally independent of tobacco given the AMR ($T \perp I \mid A$, i.e., $\alpha_1$ not significant at $P - value < 0.15$), supporting the direction $A \to I$ in both graphs. This integrated approach enables robust causal inference while accounting for the complex, multi-pathway nature of smoking's effects on PDAC prognosis. In total, 29 AMR-immune pairs supported Graph 1 and 2 pairs supported Graph 2, likely reflecting insufficient statistical power to detect real Graph 1 associations.

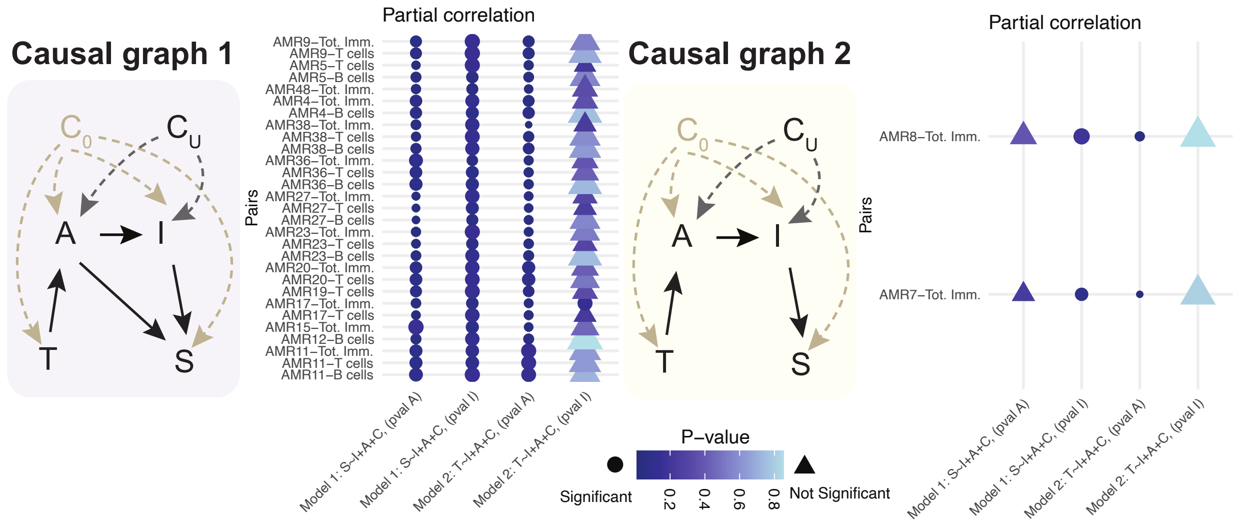


**Supplementary Figure 16: Causal discovery results showing two candidate DAG structures. Graph 1 represents pathways where AMRs have effects on both immune infiltration and survival. Graph 2 shows pathways where AMRs affect survival solely through immune infiltration. Heatmaps display partial correlation analysis results, where each point represents an AMR-immune pair and indicates the significance of associations in conditional independence models used for causal graph discrimination. Color indicates the p-value (darker = more significant), and shape distinguishes statistically significant results (pval $\leq$ 0.15). Only significant immune–AMR pairs meeting selection criteria (DAG1 or DAG2) are shown. See Methods for model details.**

##### 4.3 Serial mediation analysis of AMR-Immune pathways

Subsequently, we employed a more liberal $P - value$ threshold (0.15) for the remaining causal discovery steps to maximize sensitivity for potential associations, anticipating that only a subset of AMR-immune pairs would advance to serial mediation validation. Final candidate selection relied on 90% confidence intervals, identifying nine significant AMR-immune pairs. All models were adjusted for age, sex, and tumor grade.

Next, we sought to identify whether indirect effects exist between tobacco exposure and PDAC patient survival through identified mediator chains (AMR-immune infiltration pairs identified by causal discovery), focusing our work on pairs supported by Graph 1 (see Figure 16). In the context of multiple mediators, it is possible to decompose the total effect by considering potential mediators simultaneously, with several decomposition approaches available (26) . We adopted a counterfactual approach, adapting a recent R implementation proposed by Zugna et al. (27) to parametric survival models. We selected the "Extended imputation-based approach, estimation procedure and bootstrap without interaction between A and M1 and M2 for conditional effects" because it is the only implementation that allows decomposition of effects mediated by the first mediator (AMR, $M_1$) and the second mediator (immune infiltration, $M_2$). This sequential method first estimates the effect mediated by $M_1$, then estimates the joint mediated effect through both $M_1$ and $M_2$. The partial indirect effect via $M_2$ without passing through $M_1$ is then estimated using the extended imputation approach (Supplementary Figure 17A). For each pair of AMR–immune, latent factors simultaneously correlated with both AMR and immune variable (Section **??**) were included as additional covariates in the immune-mediator regression; when no latent factor met this criterion the original specification (inclusion of observed confounder age, gender and grade) was retained. We applied this method to all pairs identified as following the sequential pathway $T \to M_1 \to M_2 \to S$ in our causal discovery framework (Supplementary Figure 17B). As expected, we observed no indirect effects via $M_2$ alone, since immune infiltration was not directly dependent on tobacco exposure in our dataset.

We analysed serial mediation between selected AMR pairs and immune-cell–type proportions by adapting the extended imputation approach of Zugna et al. (27) to accommodate time-to-event outcomes.

Guided by the DAG validated in the previous section, we decomposed the effect of tobacco on survival into direct, methylation-mediated, and immune-mediated components. We adopted an extended imputation strategy without inverse-probability weighting, which is equivalent to the sequential plug-in $g$-formula described by Daniel et al. (26) but avoids the variance inflation associated with mediator weights. We chose a non-weighted approach because we performed extensive causal discovery before running serial mediation analysis. Therefore, we made the assumption that we identified the entire set of possible causal chains, with no back-door path—i.e., no alternative route linking tobacco to the mediators through a shared confounder (28).

**Adjustment for latent factors.** To account for unmeasured confounding detected in the *latent-factor assessment*, we refined the serial-mediation pipeline as follows: for each selected AMR–immune pair, latent factors that are significantly correlated with both the AMR and the immune profile are added as covariates in the immune-mediator regression (regression $I$). When no such latent factor is identified, regression $I$ keeps its original specification.

**Step 1: Outcome model.** For each AMR/immune pair we fitted an accelerated-failure-time model

$$\log S = \gamma_0 + \gamma_T\, T + \gamma_A\, A + \gamma_I\, I + \boldsymbol{\gamma}_C^{\top}\mathbf{C} + \varepsilon, \qquad \varepsilon \sim \mathcal{N}(0, \sigma^2), \tag{6}$$

selecting the baseline distribution (Weibull, Gaussian, log-normal, logistic, log-logistic, or exponential) by minimizing the AIC.

**Step 2: Serial imputation.** Each patient was copied four times to represent the counter-factual scenarios required by a two-step mediation chain. Counter-factual values of methylation ($A$) and immune infiltration ($I$) were imputed from linear models

$$A = \beta_0 + \beta_T\, T + \boldsymbol{\beta}_C^{\top}\mathbf{C} + \eta_A,$$

$$\text{Regression } I: \quad I = \alpha_0 + \alpha_T\, T + \alpha_A\, A + \begin{cases} \boldsymbol{\alpha}_C^\top \mathbf{C} + \eta_I, & \text{(no LF)} \\ \boldsymbol{\alpha}_U^\top \mathbf{U}^* + \boldsymbol{\alpha}_C^\top \mathbf{C} + \eta_I, & \text{(with LF)} \end{cases}$$

where $\mathbf{U}^*$ is the subset of latent factors simultaneously correlated with the AMR and the immune variable under study.

**Step 3: Effect decomposition.** The imputed survival times were regressed in the indicator variables $(a_3, a_1, a_2)$ representing tobacco status at each stage of the counter-factual sequence, yielding the interventional analog components:

$$\text{IA–DE} = \theta_{a_3}, \qquad \text{IA–IE}_A = \theta_{a_1}, \qquad \text{IA–IE}_I = \theta_{a_2}, \qquad \text{IA–TE} = \theta_{a_3} + \theta_{a_1} + \theta_{a_2}.$$

These coefficients are interpreted as time-ratio differences on the AFT scale. The interventional analog effect answers how much longer patients would live if they all stopped smoking while keeping the methylation and immune profiles they naturally exhibit as non-smokers.

**Uncertainty.** Standard errors and 90 % percentile confidence intervals were computed from *1 000* bootstrap samples resampled at the patient level.

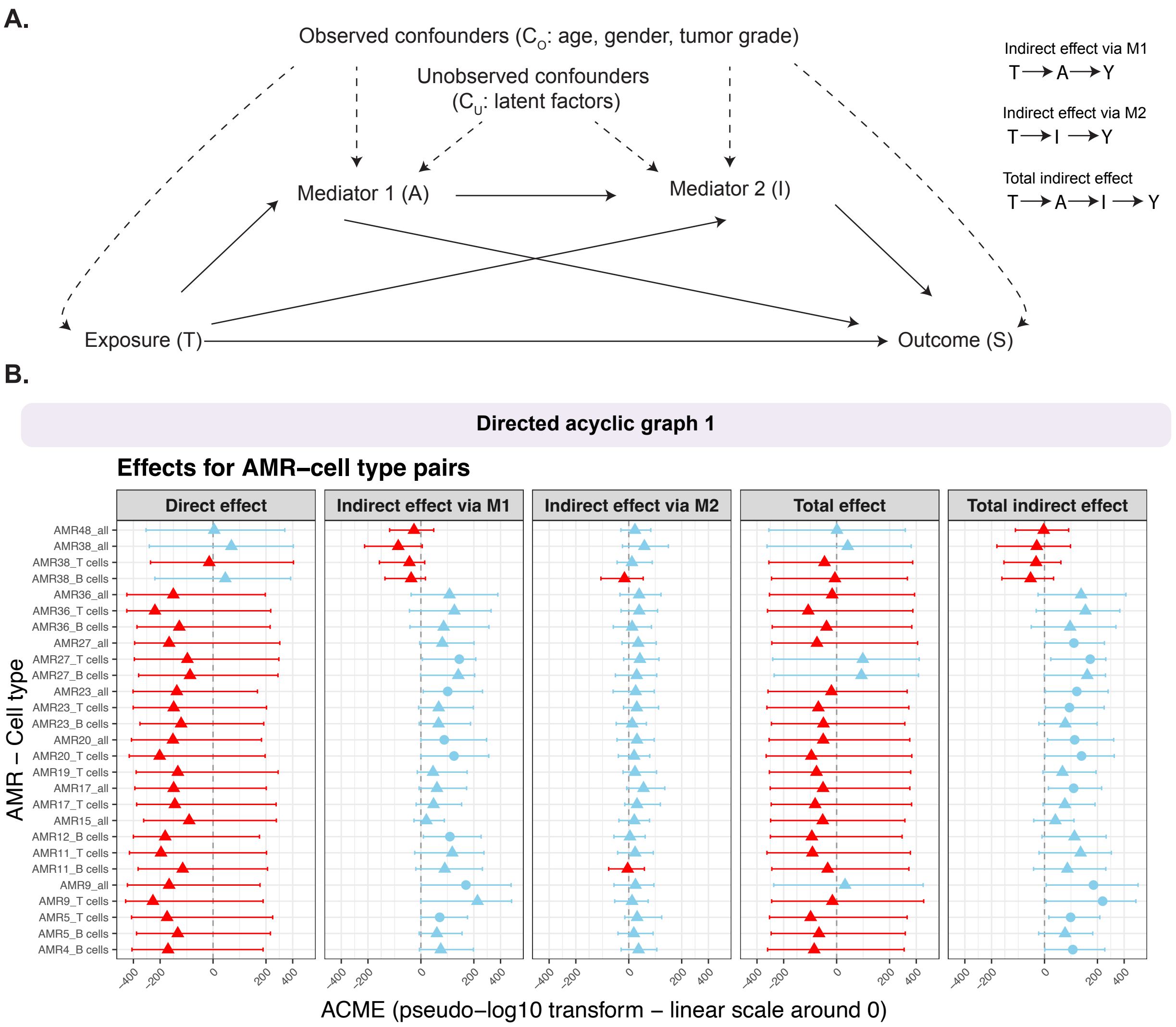


**Supplementary Figure 17: Estimates of total, direct, and indirect effects of smoking on PDAC patient survival through DNA methylation and immune infiltration. (A). DAG representing the studied structure. (B) Estimated effects for AMR-Immune pairs identified in DAG1. Circles represent significant effect, Triangle non significant effect, Colours represent blue for protective effects and red for deleterious effect. Each line represents 90 % percentile confidence intervals.**

#### 4.4 Characterization of protective and deleterious mediation pathways

We applied serial mediation analysis to the 29 pairs retained from the causal discovery step. Effect estimation failed for two pairs (AMR11–Total Immune and AMR4–B cells), leaving 27 pairs for testing. Among these, eleven pairs showed statistically significant total indirect effects, which corresponds to seven AMRs of interest (AMR4, AMR5, AMR9, AMR17, AMR20, AMR23, and AMR27). As *KCNQ1*-AMR5 and *PIWIL1*-AMR17 are already known candidates, we focused on the four AMRs containing more than two CpGs as particularly relevant examples (Figure 17B and Supplementary Figure 18). In subsequent analyses, we used refined smoking categories: never-smoker, former smoker ($>$ 15 years), former smoker ($\leq$ 15 years), and current smoker. Although more informative, these subcategories were not used in the mediation analysis to preserve statistical power by pooling into larger smoker vs. non-smoker groups. Remarkably, for all four selected AMRs (*ZBTB42*-AMR4, *CLGN*-AMR9, *RAD51AP2*-AMR20, and *PRDM16*-AMR27) we observed an indirect protective effect of tobacco exposure on survival (Figure 17B).

For *ZBTB42*-AMR4, the effect was mediated jointly by B cells. AMR4 spans the coding region of *ZBTB42*, a zinc finger protein family member not previously linked to PDAC. Tobacco exposure induced slight demethylation of this constitutively methylated locus, particularly in current smokers (Supplementary Figure 18A). Interestingly, *ZBTB42* has recently been identified as a prognostic factor associated with immune cell infiltration in glioma (29). For *CLGN*-AMR9, the effect was mediated jointly by T cells. AMR9 lies in a CpG-dense promoter region of Calmegin, a testis-specific endoplasmic reticulum chaperone. While the locus is generally unmethylated, partial methylation was observed in some smoking patients (Supplementary Figure 18B). Although not previously linked to PDAC, Calmegin has been proposed as a predictor of clear cell renal carcinoma (30) and is differentially methylated in metabolic dysfunction–associated steatotic liver disease (31). For *RAD51AP2*-AMR20 and *PRDM16*-AMR27, tobacco exposure also demonstrated protective indirect effects, mediated either by the AMR alone or jointly with T-cell infiltration (Supplementary Figure **??**C–D). AMR20 is located in the promoter of RAD51AP2, which is constitutively unmethylated but shows partial methylation in smokers. RAD51AP2 interacts with RAD51, a key DNA repair factor. While RAD51AP2 has been implicated in cholangiocarcinoma progression (32), its role in PDAC has not, to our knowledge, been previously reported. Finally, AMR27 is located in the gene body of *PRDM16*, a zinc finger transcription factor known for recurrent translocations in acute myeloid leukemia (33), but with no previously reported association with PDAC. The locus is normally hemi-methylated, yet displays a bias toward demethylation in current smokers. Importantly, across all four genes, smoking status was not associated with significant changes in gene expression (Wilcoxon rank-sum test; Supplementary Figure 18). These findings demonstrate the potential of integrating high-dimensional mediation analysis with causal discovery frameworks to uncover complex, multi-pathway mechanisms linking environmental exposures to cancer outcomes. Notably, no specific trend emerged for immune infiltration, consistent with the complexity and incomplete understanding of its role in PDAC (34). Taken together, our results highlight the intricate interplay between tobacco-induced epigenetic modifications and immune microenvironment alterations in determining PDAC patient prognosis.

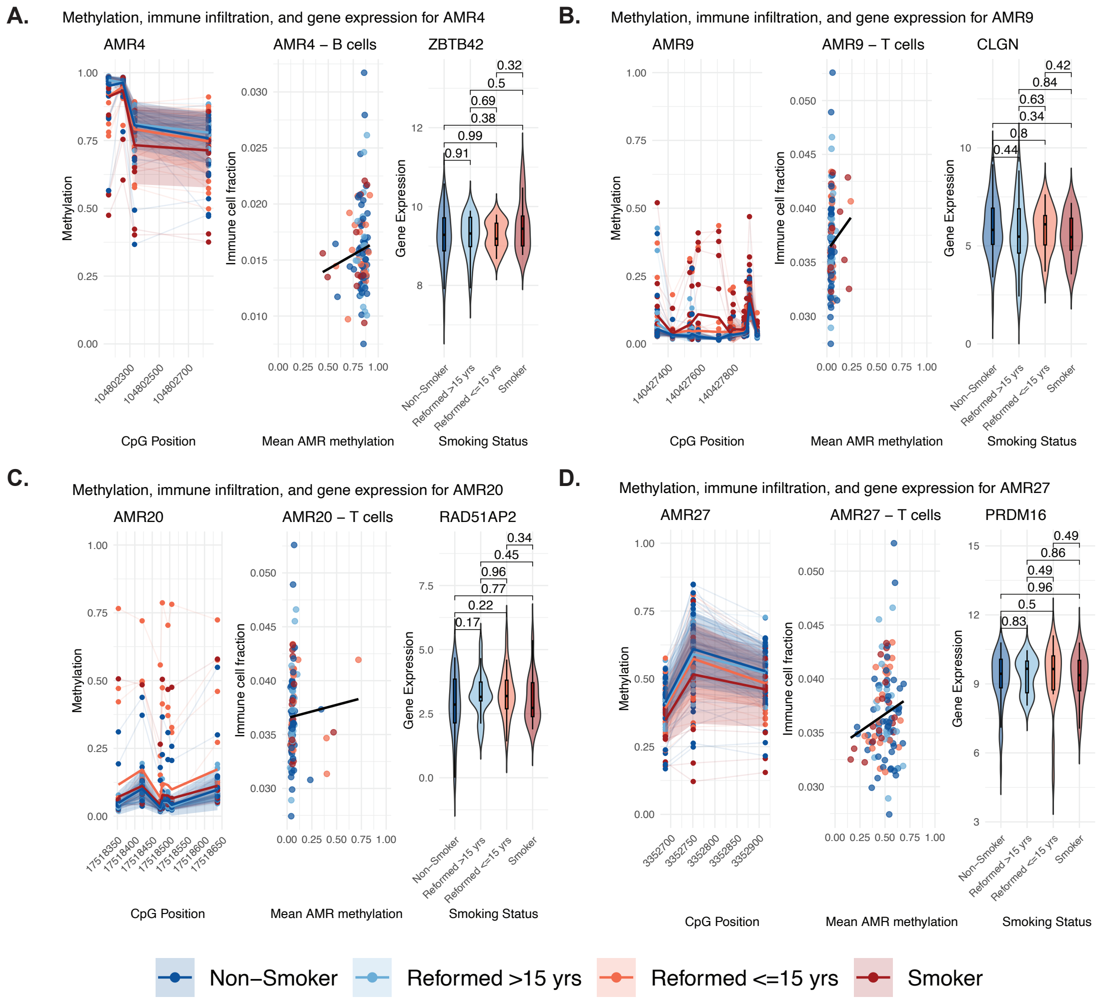


**Supplementary Figure 18: DNA methylation, immune infiltration, and gene expression associated with selected AMRs. A. *ZBTB42*-AMR4, B. *CLGN*-AMR9, C. *RAD51AP2*-AMR20, and D. *PRDM16*-AMR27. For each AMR, three panels are shown: (i) CpG site–specific methylation patterns stratified by smoking status, with individual patient trajectories (light lines), group means (solid lines), and standard deviation intervals (shaded ribbons); (ii) correlation between mean AMR methylation and immune cell fraction for the relevant cell type; and (iii) gene expression levels stratified by smoking categories; Violin plots show the distribution; boxplots indicate the median and interquartile range. P-values are from pair-wise Wilcoxon rank-sum tests. Colors are consistent across panels: red for smokers and blue for non-smokers.**

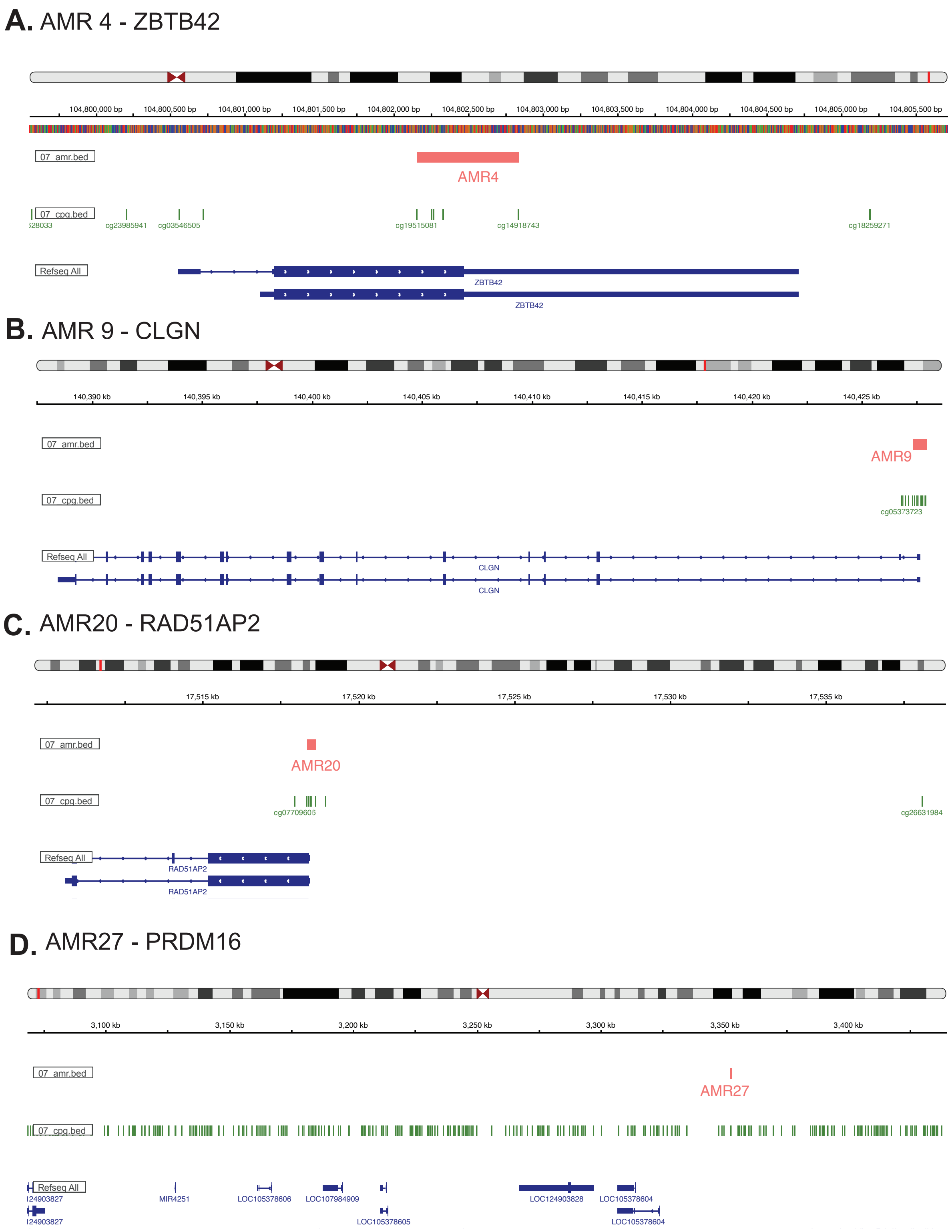


**Supplementary Figure 19:** IGV representation of the locus of know genes proximal to AMRs of interest.

### References

[1] L. Huang, *et al.*, *Bioinformatics* (2023).

[2] S. Ahn, *et al.*, *ArXiv* (2025).

[3] P. Tian, *et al.*, *Bioinformatics* (2022).

[4] A. Domingo-Relloso, *et al.*, *medRxiv: The Preprint Server for Health Sciences* (2024).

[5] B. Jumentier, *et al.*, *Environmental Health Perspectives* .

[6] D. Tingley, *et al.*, *Journal of Statistical Software* (2014).

[7] A. Jérolon, *et al.*, *The International Journal of Biostatistics* (2021).

[8] R. Nicolle, *et al.*, *Cancers* (2019).

[9] R. B. Cattell, *Multivariate Behavioral Research* (1966).

[10] K. Caye, *et al.*, *Molecular Biology and Evolution* (2019).

[11] Z. Xu, *et al.*, *Nucleic Acids Research* (2016).

[12] B. S. Pedersen, *et al.*, *Bioinformatics* (2012).

[13] S. Hamada, *et al.*, *Journal of Cellular Physiology* (2012).

[14] G. Duarte-Medrano, *et al.*, *Medicine* (2019).

[15] H. Wang, *et al.*, *Translational Cancer Research* (2020).

[16] R. Xue, *et al.*, *Digestive Diseases and Sciences* (2023).

[17] J. Xie, *et al.*, *Human Cell* (2021).

[18] F. Volta, *et al.*, *Nature Communications* (2019).

[19] J. Peng, *et al.*, *Cell Research* (2019).

[20] S. Raghavan, *et al.*, *Cell* (2021).

[21] N. U. Rashid, *et al.*, *Clinical cancer research : an official journal of the American Association for Cancer Research* (2020).

[22] M. Dong, *et al.*, *Briefings in Bioinformatics* (2021).

[23] M. Hu, *et al.*, *Bioinformatics* (2024).

[24] X. Wang, *et al.*, *Nature Communications* (2019).

[25] S. C. Zheng, *et al.*, *Nature Methods* (2018).

[26] R. M. Daniel, *et al.*, *Biometrics* (2015).

[27] D. Zugna, *et al.*, *BMC Medical Research Methodology* (2022).

[28] S. Vansteelandt, *et al.*, *Epidemiology (Cambridge, Mass.)* (2017).

[29] Y. Li, *et al.*, *Frontiers in Pharmacology* (2023).

[30] M. Cai, *et al.*, *Stem Cells International* (2023).

[31] J. Li, *et al.*, *Liver International: Official Journal of the International Association for the Study of the Liver* (2025).

[32] Q. Yao, *et al.*, *Biomedicines* (2023).

[33] I. Nishikata, *et al.*, *Blood* (2003).

[34] K. K. Mahadevan, *et al.*, *Science* (2024).
